# Reproducible Human Reward Imaging Phenotypes Exhibit Differential Sensitivity to Dopamine D2 Receptor Antagonism

**DOI:** 10.64898/2026.05.14.724267

**Authors:** Nicola Sambuco, Antonella Lupo, Peter C.T. Hawkins, Pierluigi Selvaggi, Linda A. Antonucci, Alessandro Bertolino, Giuseppe Blasi, Piergiuseppe Di Palo, Luigi Grassi, Daniela Grasso, Philipp Homan, Gian Marco Leggio, Francesco Massari, Alessio Maria Monteleone, Martin Osugo, Roberta Passiatore, Alessandra Raio, Antonio Rampino, Tobias Banaschewski, Gareth J. Barker, Arun L.W. Bokde, Rüdiger Brühl, Sylvane Desrivières, Herta Flor, Hugh Garavan, Penny Gowland, Antoine Grigis, Andreas Heinz, Jean-Luc Martinot, Marie-Laure Paillère Martinot, Eric Artiges, Frauke Nees, Dimitri Papadopoulos Orfanos, Luise Poustka, Michael N. Smolka, Nathalie Holz, Nilakshi Vaidya, Henrik Walter, Robert Whelan, Gunter Schumann, Apulian Network on Risk for Psychosis, Oliver D. Howes, Mitul A. Mehta, Giulio Pergola

**Affiliations:** Department of Translational Biomedicine and Neuroscience, University of Bari Aldo Moro, 70124 Bari, Italy; Department of Neuroimaging, Institute of Psychiatry, Psychology and Neuroscience, King’s College London, WC2R 2LS London, UK; Psychiatric Unit - University Hospital, 70124 Bari, Italy; Department of Neuroscience and Rehabilitation, Institute of Psychiatry, University of Ferrara, Ferrara, Italy; Department of Neuroradiology, IRCCS Casa Sollievo della Sofferenza, 71013 San Giovanni Rotondo, Foggia, Italy; Psychiatric Hospital, University of Zurich, Switzerland; Neuroscience Center Zurich, University of Zurich & Swiss Federal Institute of Technology Zurich, Zurich, Switzerland; Department of Biomedical and Biotechnological Sciences, University of Catania, 95123, Catania, Italy; Department of Psychiatry, University of Campania “Luigi Vanvitelli”, Naples, Italy; Department of Psychosis Studies, Institute of Psychiatry, Psychology & Neuroscience, King’s College London, London, UK; MRC Laboratory of Medical Sciences, Imperial College London, London, UK; South London and Maudsley NHS Foundation Trust, London, UK; Lieber Institute for Brain Development, Johns Hopkins Medical Campus, 21205 Baltimore, MD, United States; Department of Child and Adolescent Psychiatry and Psychotherapy, Central Institute of Mental Health, Medical Faculty Mannheim, Heidelberg University, Square J5, 68159 Mannheim, Germany; German Center for Mental Health (DZPG), partner site Mannheim-Heidelberg-Ulm; Discipline of Psychiatry, School of Medicine and Trinity College Institute of Neuroscience, Trinity College Dublin, Dublin, Ireland; Physikalisch-Technische Bundesanstalt (PTB), Braunschweig and Berlin, Germany; Social, Genetic and Developmental Psychiatry Centre, Institute of Psychiatry, Psychology & Neuroscience, King’s College London, United Kingdom; Institute of Cognitive and Clinical Neuroscience, Central Institute of Mental Health, Medical Faculty Mannheim, Heidelberg University, Square J5, Mannheim, Germany; Department of Psychology, School of Social Sciences, University of Mannheim, 68131 Mannheim, Germany; Departments of Psychiatry and Psychology, University of Vermont, 05405 Burlington, Vermont, USA; Sir Peter Mansfield Imaging Centre School of Physics and Astronomy, University of Nottingham, University Park, Nottingham, United Kingdom; NeuroSpin, CEA, Université Paris-Saclay, F-91191 Gif-sur-Yvette, France; Department of Psychiatry and Psychotherapy, University of Tübingen, Germany; German Center for Mental Health (DZPG), Site Tübingen, Germany; Institut National de la Santé et de la Recherce Médicale, INSERM U A10 “Trajectoires développementales & psychiatrie”, University Paris-Saclay, Ecole Normale Supérieure Paris-Saclay, CNRS; Centre Borelli, Gif-sur-Yvette, France; AP-HP. Sorbonne Université, Department of Child and Adolescent Psychiatry, Pitié-Salpêtrière Hospital, Paris, France; Psychiatry Department, EPS Barthélémy Durand, Etampes, France; Institute of Medical Psychology and Medical Sociology, University Medical Center Schleswig Holstein, Kiel University, Kiel, Germany; Department of Child and Adolescent Psychiatry, Center for Psychosocial Medicine, University Hospital Heidelberg, Heidelberg, Germany; Department of Psychiatry and Psychotherapy, Technische Universität Dresden, Dresden, Germany; Centre for Population Neuroscience and Stratified Medicine (PONS), Department of Psychiatry and Psychotherapy, Charité Universitätsmedizin Berlin, Germany; Department of Psychiatry and Psychotherapy CCM, Charité – Universitätsmedizin Berlin, corporate member of Freie Universität Berlin, Humboldt-Universität zu Berlin, and Berlin Institute of Health, Berlin, Germany; School of Psychology and Global Brain Health Institute, Trinity College Dublin, Ireland; Centre for Population Neuroscience and Precision Medicine (PONS), Institute for Science and Technology of Brain-inspired Intelligence (ISTBI), Fudan University, Shanghai, China; Department of Mental Health, ASL Foggia, Foggia, IT; Department of Clinical and Experimental Medicine, University of Foggia, Foggia, IT; Department of Mental Health, ASL Barletta-Andria-Trani, Andria, IT; Department of Mental Health, ASL Bari, Bari, IT; Department of Mental Health, ASL Brindisi, Brindisi, IT; Department of Psychiatry and Behavioral Sciences, Johns Hopkins University School of Medicine, 21205 Baltimore, MD, United States; Department of Genetic Medicine, Johns Hopkins University School of Medicine, 21205 Baltimore, MD, United States

**Author notes:** **Corresponding authors:** Giulio Pergola, Address: Piazza Giulio Cesare, 11 - 70124 Bari, Italy –, Mitul Mehta, Address: Denmark Hill, London, SE5 8AF - England. These authors contributed equally to this work and share the first authorship (Nicola Sambuco, Antonella Lupo).

## Abstract

Human reward processing varies along cue-centric and outcome-centric axes, but a reproducible mechanistic account of individual variation in incentive salience attribution has been lacking. Using fMRI across five cohorts (N-total=1,251; N1=890; N2=245; N3=34; N4=48; N5=34), we identified two robust imaging phenotypes mirroring sign- and goal-tracking (ST-like, GT-like). ST-like individuals showed dominant ventral striatal responses to reward-anticipation cues and sustained incentive salience attribution; GT-like individuals showed heightened responses to reward outcomes. This distinction was replicable across sites and independent samples. Single-dose and repeated-dose D_2_/D_3_ antagonism (risperidone, haloperidol, amisulpride) selectively reduced anticipatory ventral striatal activity in ST, with single-dose antagonism additionally producing a parallel drop in self-reported energy. Instead, D_2_/D_3_ partial agonism (aripiprazole) increased anticipatory and reduced outcome-phase responses in GT. In a psychosis cohort, antipsychotic D_2_ affinity was associated with blunted anticipatory signals and higher negative symptom burden, offering a neuroimaging-driven basis for stratifying patients and predicting response to dopaminergic agents.

## INTRODUCTION

Alterations in reward processing are a fundamental component of psychiatric disorders^1–5^, and the study of reward brain circuits has spurred investigations that, in humans, primarily focus on two distinct temporal phases: the anticipation phase, when cues trigger reward prediction, and the outcome phase, when feedback is evaluated^6^. Building on these human findings, the Sign-Tracker/Goal-Tracker (ST/GT) model, derived from rodent studies, complements this phase-based approach by emphasizing individual differences in cue-reward learning and integrating both the reward anticipation and outcome phases^7,8^. Indeed, rodent research has shown increased striatal activity in ST versus GT in response to reward-associated cues, reflecting excessive incentive salience attribution and reliance on dopamine-mediated reward prediction errors^7–10^. By contrast, GT focuses on the reward outcome, demonstrating a more cognitively controlled strategy that facilitates goal-directed behavior^11^. A third subset of animals, termed intermediate, exhibits both sign- and goal-tracking conditioned responses at different trials with no clear preference for either; in the present work, primary analyses focus on the ST-like and GT-like profiles, with intermediates considered in additional analyses. Yet only a few attempts have been made to replicate this in humans, relying on small samples and complex learning paradigms^12–15^ that are poorly suited for clinical research, particularly among populations with cognitive and learning deficits^16–18^. Because humans in real-world contexts primarily interact with cues that already carry reward associations^19^, the neural substrates of human ST/GT-like behavior remain largely unexplored.

In humans, ST-like behavior manifests as automatic attentional capture by reward-predicting cues, whereas GT-like behavior is characterized by a preferential orientation toward the reward-delivery context^13,15,20^. Outside controlled learning paradigms, this dissociation co-segregates with individual differences in impulsivity, compulsivity, and addiction-prone tendencies, measurable even in daily-life contexts using ecological momentary assessment^21^. These traits are transdiagnostically relevant across psychiatric conditions that share disrupted cue-driven motivation. Of relevance for psychiatric populations, schizophrenia is characterized by circuit-specific dysregulation of dopaminergic signaling, in which aberrant attribution of incentive salience to irrelevant stimuli coexists with blunted ventral striatal responses to reward-predictive information and maps onto the positive and negative symptom dimensions of the disorder^22^. This pattern raises the possibility that individual differences in cue- versus outcome-driven reward processing modulate striatal responsiveness to D_2_/D_3_-targeting antipsychotics and shape symptom presentation in schizophrenia.

The first aim of the current study was to assess whether ST/GT phenotypes emerge in humans on a task where cue-reward contingencies are instructed in advance, while still enabling distinct evaluation of reward anticipation and outcome. Recognizing that behavior fundamentally arises from brain activity^23–26^, here we focused on the neural substrates of reward processing. We used functional magnetic resonance imaging (fMRI) to examine brain responses during the Monetary Incentive Delay (MID) task^27^. In the MID task, cue-reward associations are explicitly instructed during a pre-scan training session, separating learning from task performance. Brain activity is then measured as cues signal potential monetary reward (anticipation) and as participants receive performance-based feedback on a simple discrimination task (outcome). Unlike the Pavlovian conditioned approach paradigm used in rodents, the MID task does not require new cue-reward learning during scanning or measure overt approach behavior but preserves the key translational feature of dissociating neural responses to reward-predicting cues from those to reward outcomes, the functional basis for distinguishing ST-like from GT-like phenotypes.

In our Discovery cohort, we utilized data from an open-access consortium of 890 healthy young adults to identify two distinct neurocognitive profiles analogous to the ST/GT phenotypes observed in rodent models. We hypothesized that individuals with ST tendencies would exhibit increased neural activity during reward anticipation, whereas those with GT tendencies would show greater activation during reward outcomes. Given the heightened sensitivity of ST to incentive cues^28^, we predicted that ST would show greater risk adjustment on the Cambridge Gambling Task (CGT) and more reward-driven responding on the Passive Avoidance Learning Paradigm (PALP). Given rodent evidence that ST rats outperform GT on set-shifting, we predicted superior executive flexibility in human ST on the Wisconsin Card Sorting Test (WCST). Personality predictions were exploratory, given the limited prior human evidence linking ST/GT-like behavior to personality. To validate these findings, we analyzed an independent sample of 245 healthy participants who completed a different version of the MID task and additional neuropsychological assessments. Because rodent ST/GT differences hinge on the persistence of incentive salience across trials, we leveraged the large trial counts in the replication dataset to test whether sustained incentive salience to reward-anticipation cues differentiates ST-like from GT-like individuals^13,29^.

Once the consistency and the replicability of this neurocognitive model were confirmed, the second aim of the study was to test whether ST/GT-like neuroimaging phenotypes would predict differential responses to dopaminergic agents, motivated by preclinical evidence linking dopamine signaling to cue-driven behavior^30–32^. In rodents, ST and GT exhibit distinct sensitivities to D_2_ modulation, with cue-reactivity in ST relying more heavily on dopamine signaling than GT^10,33–35^. To address this translational question, we conducted a single-dose pharmacological study in healthy adults using a double-blind, placebo-controlled, cross-over design, comparing risperidone or haloperidol with placebo during the MID task^36,37^. We hypothesized that, in this single-dose context, ST would show selective reductions in cue-driven BOLD activity under D_2_ blockade, offering a potential mechanistic basis for targeted therapeutic strategies in disorders involving maladaptive reward processing.

However, the single-dose study left two questions open: i) whether ST-specific pharmacological sensitivity generalizes beyond acute exposure, ii) whether it generalizes across D_2_/D_3_ agents with distinct pharmacodynamics. Full D_2_/D_3_ receptor antagonism suppresses both tonic and phasic dopamine signaling, whereas partial D_2_/D_3_ agonism preferentially attenuates phasic over tonic dopamine release^38^, making the contrast between the two pharmacological classes informative for isolating the dopamine component on which ST cue-reactivity depends. To address both questions, we applied the ST/GT classification framework to an independent pharmacological neuroimaging dataset in which healthy volunteers received repeated-dose amisulpride (a selective D_2_/D_3_ antagonist) or aripiprazole (a D_2_/D_3_ partial agonist) for 7 consecutive days in a randomized, double-blind, placebo-controlled crossover design^39^. We hypothesized that repeated-dose D_2_/D_3_ antagonism with amisulpride would replicate the ST-specific attenuation of anticipatory ventral striatal BOLD observed under single-dose risperidone and haloperidol, and that the mechanistically distinct profile of aripiprazole would yield a differential pattern of phenotype-specific effects.

Building on the circuit-specific dopaminergic dysregulation outlined above, the third aim brought the ST/GT framework into a schizophrenia population to test whether the cue- versus outcome-driven dissociation maps onto interindividual variability in symptom burden and treatment response. This aim also extended the pharmacological logic developed in healthy volunteers to a clinical context, where patients receiving antipsychotic treatment accumulate D_2_/D_3_ engagement spanning weeks in first-episode psychosis (FEP) to months and years in chronic schizophrenia. Prior studies have consistently reported a blunted ventral striatal response to both reward anticipation and outcome in schizophrenia^40^, reflecting a global attenuation of incentive salience attributable, at least in part, to antipsychotic treatment^36,41^. We therefore asked whether, despite this overall attenuation, meaningful interindividual variability in cue- versus outcome-driven ventral striatal responses analogous to ST and GT profiles, operationalized as replication of the ST/GT dissociation in whole-brain voxel-wise maps and ventral striatal BOLD contrasts, could still be detected in a combined clinical sample of medicated patients with chronic schizophrenia and FEP. We predicted that ST-like patients would retain greater residual anticipatory ventral striatal activity than GT-like patients; that GT-like patients would show higher negative symptom burden given their blunted cue-driven profile; and that, among patients with documented ongoing antipsychotic treatment, stronger D_2_/D_3_ blockade would be associated with a GT-like phenotype and greater negative symptom severity^39^.

## RESULTS

### Discovery cohort

#### Cluster development

The Discovery cohort comprised 890 healthy participants from IMAGEN^42^ (454 females, 436 males; mean age=22.1 years, SD=.7 years), a multi-site European study (Figure 1, panel a). Demographic details, grouped by site, are presented in Supplementary Table 1. Participants performed the MID task during fMRI, in which a cue signaling potential reward (anticipation) was followed by a target response and feedback (outcome; Figure 1, panel b). From these data, we extracted BOLD signals from meta-analytic regions of interest spanning both phases (Figure 1, panel c; full list of ROIs in Supplementary Fig. 1)^6^. All measures reflect reward-versus-control contrasts. We then applied hierarchical k-means clustering (k=3) with Ward’s linkage to derive Sign-Tracker (ST), Goal-Tracker (GT), and Intermediate (INT) reward-related phenotypes in the Discovery cohort (Figure 1, panel d). The number of clusters was set on hypothesis-driven grounds; the data-driven analyses are reported in the Methods and Supplementary Methods (Supplementary Figs. 2-3). Modeling the intermediate group as a separate cluster isolates the ST and GT centroids on their more prototypical members, sharpening the templates subsequently projected onto the other cohorts.

**Figure 1.**
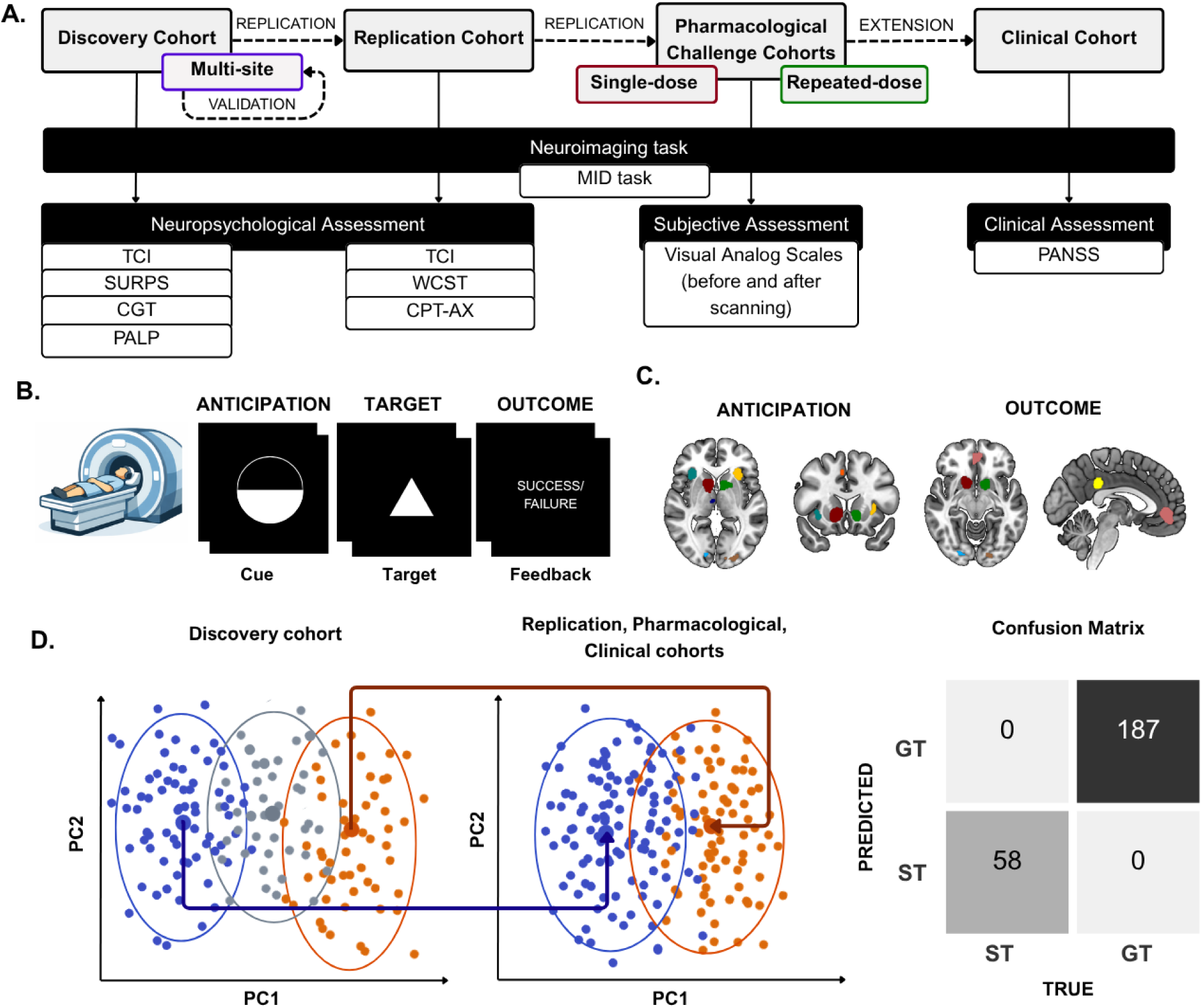
Experimental Design, fMRI Task, and Machine Learning Pipeline. **a. Study Flowchart:** Overview of the multi-stage study, including the multi-site Discovery cohort, Replication cohort, Pharmacological challenge cohorts (single-dose and repeated-dose), and Clinical cohort. Assessments include neuroimaging (MID task) across all four cohorts, neuropsychological batteries (i.e., TCI, SURPS, CGT, PALP, WCST, CPT-AX) in the Discovery and Replication cohorts, subjective assessment (VAS) in the single-dose Pharmacological Challenge cohort, and clinical scale (PANSS) in the Clinical cohort. **b. Monetary Incentive Delay (MID) Task:** Generic representation of the MID task, in which participants viewed a cue (Anticipation), responded to a target, and received feedback (Outcome) during fMRI scanning. **c. ROI-selection:** Meta-analytically derived regions of interest (ROIs) were defined for reward anticipation (bilateral ventral striatum, thalamus, supplementary motor area, bilateral motor cortex, bilateral anterior insula, bilateral lateral occipital cortex, and bilateral middle frontal gyrus) and outcome (bilateral ventral striatum, ventromedial prefrontal cortex, bilateral lateral occipital cortex, posterior cingulate cortex, bilateral superior parietal lobule, left middle frontal gyrus, and right inferior frontal gyrus). **d. Machine learning pipeline:** Clustering algorithm was first applied to the Discovery sample, identifying three data-driven clusters (left). To assess the generalizability of the clustering, the Discovery algorithm was applied to independent datasets, i.e., the Replication, Pharmacological Challenge, and Clinical cohorts, by projecting only the ST/GT centroids (center). The Confusion Matrix (right) evaluates the consistency and generalizability of cluster assignments (ST vs. GT) across datasets by comparing the predicted cluster assignments with those obtained from an independent hierarchical k-means clustering applied to the external datasets.

Cluster stability was assessed via leave-site-out validation, yielding balanced accuracy metrics for each held-out site. To this end, the dataset was initially divided into a training set (seven sites) and a testing set (one site). Scaling (range: -1 to 1) was applied separately to both sets, and residuals of regression models for the site effect were included only in the training set. Hierarchical k-means clustering was performed on the training set with k=3, using Euclidean distance and Ward’s linkage, yielding centroids for ST, GT, and the intermediate clusters. To validate this algorithm on the testing set, each held-out participant was assigned to a cluster by computing the Euclidean distance to the ST and GT training centroids only, as the limited size of single-site testing samples did not support reliable recovery of the smaller intermediate group; full methodological details are provided in the Supplementary Methods.

We then applied the clustering procedure to the testing set to obtain cluster labels. We calculated a confusion matrix (Figure 1, panel d) using the *caret* R package to evaluate the cross-tabulation of cluster assignments and associated statistics, i.e., sensitivity, specificity, balanced accuracy, and overall accuracy. Overall accuracy was reported with 95% confidence intervals and tested against the no-information rate, and balanced accuracy, derived from sensitivity and specificity, served as the primary replicability metric.

Figure 2, panel a, illustrates that the three-cluster solution in the IMAGEN cohort identified the following groups: ST (N=161), who exhibited greater BOLD responses during reward anticipation relative to outcome; GT (N=319), who exhibited greater BOLD responses during outcome relative to anticipation; and an intermediate group of subjects (N=410) displaying no preferential effects on either phase. No statistically significant sex differences were observed, supported by two converging analyses. First, the distribution of males and females did not differ between ST (M=89, F=72) and GT (M=149, F=170), χ^2^_(1, N=480)_=2.81, p=0.094, and second, very similar imaging phenotypes emerged when the cluster analysis was conducted separately for males and females (Supplementary Fig. 4).

**Figure 2.**
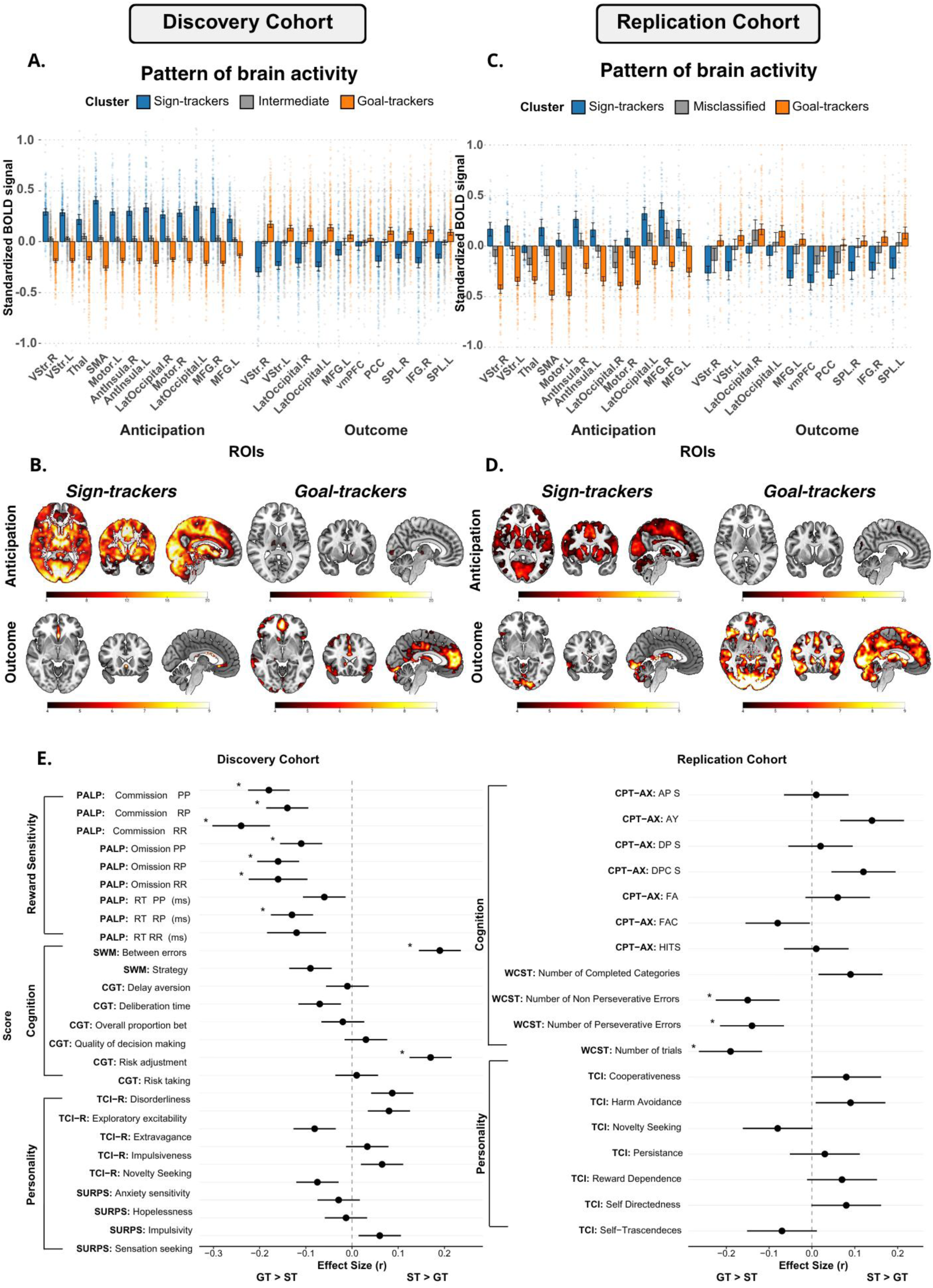
Brain activity patterns and behavioral characterization of Sign-trackers and Goal-trackers clusters. **a. Pattern of brain activity:** A three-cluster solution identified sign-trackers (ST), goal-trackers (GT), and an intermediate group. Bars represent 95% CI. **b. Voxel-wise activation maps:** Whole-brain analyses confirmed strong anticipatory BOLD activation in ST and outcome activation in GT, with minimal engagement in the alternate phase. All statistics were corrected for multiple comparisons using the Threshold-Free Cluster Enhancement (TFCE) approach ^43^, with significance set at a familywise error rate (FWE) of *_α_*<0.05. **c. Pattern of brain activity:** Generalization in a replication cohort showed high classification accuracy (82.4%), though sensitivity was reduced. **d. Voxel-wise activation maps:** Whole-brain analyses in the replication cohort replicated the key findings (p_TFCE-FWE_<0.05). **e. Cognitive and Personality Profiling:** Bars depict the effect size r of the Wilcoxon rank-sum tests performed for the associations between cluster membership (GT vs. ST) and neuropsychological measures. Left: Discovery cohort results covering Reward Sensitivity (PALP), Cognition (SWM, CGT), and Personality (TCI-R, SURPS). Right: Replication cohort results for Cognition (CPT-AX, WCST) and Personality (TCI). Positive values indicate higher scores in ST, while negative values indicate higher scores in GT (*pFDR < 0.05).

Whole-brain voxel-wise analyses of ST and GT during anticipation and outcome phases showed significant extensive activation in ST during anticipation and in GT during outcome (Figure 2, panel b), with minimal activation during the alternate phase (threshold-free cluster enhancement with family-wise error correction, pTFCE-FWE < 0.05; Supplementary Table 2). Despite inter-site variability (Supplementary Fig. 5), the leave-site-out (LSO) cross-validation analysis to assess cluster stability produced an overall accuracy of 86.7% and a measure of replicability of 84.9% (SD=9%), based on the average between sensitivity (87.8%) and specificity (82%).

#### Behavioral analyses in the MID task

Because the MID task used in the Discovery cohort implements an online tracking algorithm that adjusts target duration to maintain a 66% success rate (Supplementary Methods), hit rates are equalized across participants by design, and no phenotype-related differences in task performance were expected. Consistent with this, ST and GT hit rates converged near the 66% titration target (GT: mean=65.05, SD=12.74; ST: mean=63.4, SD=10.9; F(1, 470)=6.094, p=0.014), with higher hit rates in Reward than No-Reward trials (mean=66.92, SD=11.58 vs. mean=62.06, SD=12.27; F(1, 470)=25.59, p<.001) and no Cluster × Condition interaction (F(1, 470)=0.3, p=0.585). The 1.6-point ST/GT difference, although statistically detectable in N=472, falls within the noise of the titration algorithm and rules out differential task engagement as a driver of the BOLD-defined ST/GT distinction.

#### Personality and cognitive profiles of ST and GT

Group differences in temperament and personality were assessed using the Temperament and Character Inventory–Revised (TCI-R)^44,45^, and the Substance Use Risk Profile Scale (SURPS)^46^, while decision-making was evaluated with the Cambridge Guessing Task (CGT; https://www.cambridgecognition.com/cantab/). Passive avoidance learning was assessed using the Passive Avoidance Learning Paradigm (PALP)^47^, in which participants learn, by trial and error, to respond to rewarded stimuli and withhold responses to punished stimuli. Outcome measures include commission errors (responses on no-go trials), omission errors (nonresponses on go trials), and reaction times across reward-reward (RR), punishment-punishment (PP), and reward-punishment (RP) contingency conditions (see Methods). All group comparisons employed Wilcoxon rank-sum tests, using the False Discovery Rate (FDR)^48^ correction (α < 0.05) for each set of measures. Mean (SD) values for ST and GT, along with full statistical details of group effects, are reported in Supplementary Table 3 and Figure 2, panel e. No significant differences between ST and GT were observed in personality traits, as assessed by the TCI and SURPS.

We found a significant difference in decision-making between groups on the CGT, with ST scoring higher than GT on risk adjustment (p_FDR_=0.002), a measure of how much participants adjust their bets based on their perceived probability of winning. This pattern is consistent with the heightened sensitivity to incentive cues characterizing the ST profile. Significant differences emerged between groups for reward-related behavior (PALP). In conditions without contingency switches, GT made significantly more omission errors (nonresponses to ‘go’ trials, indicating too much behavioral inhibition) and commission errors (responses to ‘no-go’ trials, indicating too little behavioral inhibition) in both the reward-reward (Omission: p_FDR=_0.01; Commission: p_FDR=_0.002) and punishment-punishment (Omission: pFDR=0.001; Commission: p_FDR_ < 0.001) conditions. This suggests that GT individuals struggle with a lack of behavioral inhibition (commission errors) and an excess of it (omission errors), indicating a general impairment in learning-based behavioral regulation under reward and punishment contingencies. The reward-learning dissociation (PALP) was reproduced when these analyses were repeated using a k=2 solution that partitions participants directly into ST and GT, indicating that the core behavioral difference does not depend on the exclusion of the intermediate group (Supplementary Methods and Supplementary Fig. 6). The same dissociations were preserved when ST and GT were defined by a percentile criterion on the first principal component instead of by clustering. Classifying the most ST and most GT 45%, 40%, and 30% of participants as the two phenotypes, and leaving the remaining 10%, 20%, and 40% as intermediate, the core reward-learning (PALP), spatial working-memory (SWM), and risk-adjustment (CGT) differences remained significant in the same direction at every threshold (Supplementary Methods and Supplementary Fig. 7).

### Replication cohort

#### Cluster replicability

The independent Replication cohort of 245 healthy individuals was recruited at the University of Bari Aldo Moro (UNIBA; 141 females, 104 males; mean age=25.7 years, SD=5.7). Cluster replicability was quantified in this sample by applying the Discovery ST/GT centroids to the independent dataset and computing confusion-matrix metrics—overall accuracy, sensitivity, specificity, and replicability. Cluster replicability in the Replication cohort showed good clustering performance (Figure 2, panel c), with 82.4% overall accuracy, perfect specificity (100%), but lower sensitivity (57.4%), and 78.7% replicability based on sensitivity and specificity. While all GT were correctly identified, 43 of the Discovery-predicted ST were assigned to a different cluster by the independent Replication clustering (mis-ST), consistent with the intermediate phenotype observed in rodent models. The mis-ST were excluded from further analyses. Consistent with the Discovery cohort, whole-brain voxel-wise analyses (p_TFCE-FWE_ < 0.05) showed extensive ST activation during anticipation and GT activation during outcome (Figure 2, panel d), with minimal activation in the alternate phase (Supplementary Table 2).

#### Behavioral analyses in the MID task

Similar to the Discovery cohort, the MID task used in the Replication cohort implements an online tracking algorithm that calibrates target duration based on training-phase performance to maintain at least a 66% success rate (Supplementary Methods). Then, equivalent hit rates across phenotypes are an expected feature of the task design. Consistent with this, ST and GT hit rates were equivalent (mean=73.99, SD=6.33 vs. mean=73.68, SD=6.62; F(1, 198)=0.251, p=0.617), with no Reward vs. No-Reward effect (mean=75.49, SD=6.04 vs. mean=72.05, SD=6.57; F(1, 198)=0.001, p=0.974) and no Cluster × Condition interaction. Convergence of hit rates around the titration target indicates that the ST/GT BOLD dissociation reflects neurobiological variability under matched task performance, rather than differences in task engagement or task validity.

#### Personality and cognitive assessment of ST and GT

Group differences in personality (TCI)^45^ and sustained attention assessed through the Continuous Performance Test (CPT)^49^, as well as executive function via the Wisconsin Card Sorting Test (WCST)^50,51^, were computed using Wilcoxon rank-sum tests, setting significance at p_FDR_<0.05 across each set of comparisons. Mean (SD) values and full statistical details are in Supplementary Table 3. No significant differences in ST-GT were found on personality (TCI) or CPT measures of sustained attention, impulsivity, and vigilance. Regarding executive functions, in the WCST, ST used significantly fewer trials (p_FDR_ = 0.026) and made significantly fewer perseverative and non-perseverative errors (p_FDR_ = 0.042) than GT. The number of completed categories did not differ significantly between groups (p = 0.13). Results are depicted in Figure 2, panel e. Altogether, these results suggest that while ST and GT do not differ on measures of personality or sustained attention, the ST phenotype is associated with superior set-shifting performance, suggesting greater cognitive flexibility. We further examined the influence of the intermediate group by repeating the analyses using a k=2 solution, which directly partitions participants into ST and GT without an intermediate category. Under this partition, the set-shifting (WCST) difference did not reach significance, consistent with smaller effects being diluted when intermediate participants are included (Supplementary Methods and Supplementary Fig. 6).

A complementary trial-by-trial mixed-effects analysis of ventral striatal responses, testing whether the ST/GT dissociation is stable across the task, is reported in the Supplementary Methods (Supplementary Fig. 8).

### Pharmacological challenge cohorts

#### Single-dose

##### Cluster replicability

The single-dose D_2_ antagonist cohort included 34 healthy males (mean age=26.9 years, SD=6.8) from a placebo-controlled, crossover, single-dose study receiving haloperidol 3mg, risperidone 0.5mg, or risperidone 2mg^36,37^. More details on this cohort and study design are provided in the Supplementary Methods. Cluster replicability was evaluated using confusion-matrix metrics (accuracy, sensitivity, and specificity). Cluster replicability in this additional external validation cohort was high (92.3%), with 94.1% accuracy, 84.6% sensitivity, and perfect specificity (100%), misidentifying only 2 out of 13 ST (Figure 3, panel a) that were excluded from subsequent analyses. Following exclusion, the final sample comprised 11 ST and 21 GT, each receiving all drug conditions in a crossover design.

**Figure 3.**
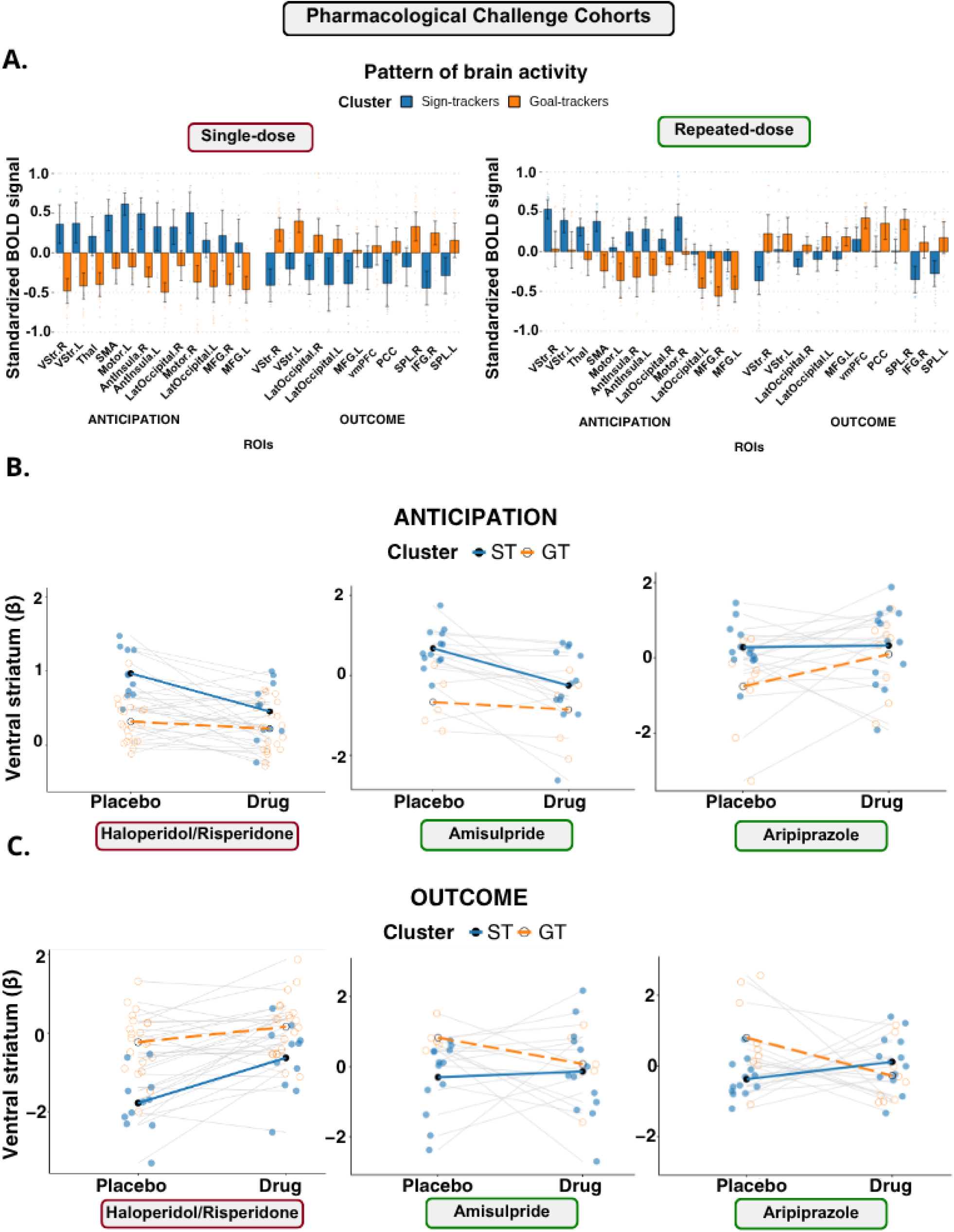
Pharmacological challenge cohorts. **a. Pattern of brain activity:** A two-cluster solution identified sign-trackers (ST) and goal-trackers (GT). Standardized BOLD signals in selected ROIs during Anticipation and Outcome phases for the *Single-dose* (left) and *Repeated-dose* (right) cohorts. The divergent activation patterns between ST (blue) and GT (orange) remain consistent across pharmacological groups. **b. Drug effects on ventral striatal reactivity during anticipation:** Comparison of Placebo vs. Drug (Haloperidol/Risperidone, Amisulpride, and Aripiprazole) on Ventral Striatum beta values (β) during Anticipation. Individual trajectories (grey lines) and cluster averages (colored lines) illustrate how different dopaminergic antagonists and partial agonists differentially modulate striatal signaling in ST and GT phenotypes, suggesting a cluster-specific sensitivity to dopamine receptor manipulation. **c. Drug effects on ventral striatal reactivity during outcome:** Comparison of Placebo vs. Drug (Haloperidol/Risperidone, Amisulpride, and Aripiprazole) on Ventral Striatum beta values (β) during Outcome phase. Individual trajectories (grey lines) and cluster averages (colored lines) illustrate how different dopaminergic antagonists and partial agonists differentially modulate striatal signaling in ST and GT, suggesting a cluster-specific sensitivity to dopamine receptor manipulation.

##### Placebo vs. Drug effect on ventral striatal reactivity

Drug effects on bilateral ventral striatum BOLD were tested using a linear mixed-effects model with Phase (anticipation, outcome), Drug condition (placebo, drug), and Cluster (ST, GT) as fixed factors, including all interactions. The model controlled for drug type (risperidone or haloperidol) and age (modeled as a second-order polynomial), and included subject as a random intercept. Results are depicted in Figure 3, panels b and c. The Phase × Drug × Cluster interaction was significant (F(1,90) = 4.65, p = .034, η²p = .049), indicating that drug effects on ST and GT differed across task phases. In the anticipation phase, the Drug × Cluster interaction was significant (F(1,30) = 11.58, p = .002, η²p = .279). Under placebo, ST showed higher BOLD responses than GT (β = 0.77, SE = 0.13, t(39.6) = 5.99, p < .001, d = 3.56), and drug significantly reduced ST responses (β = −0.41, SE = 0.09, t(30) = −4.48, p < .001, d = 1.91) with no effect in GT (β = −0.03, SE = 0.07, t(30) = −0.38, p = .706, d = 0.12). In the outcome phase, the Drug × Cluster interaction did not reach significance (F(1,30) = 2.87, p = .101, η²p = .087). Given the significant three-way interaction, planned contrasts within the outcome phase revealed a significant drug-related increase in ST (β = 0.98, SE = 0.32, t(30) = 3.11, p = .004, d = 1.33), with no significant change in GT (β = 0.32, SE = 0.23, t(30) = 1.41, p = .168, d = 0.44).

Additionally, a dose-response analysis of risperidone in the anticipation phase, conducted in the subsample receiving both dose levels (N = 17), confirmed a significant linear decrease in ST BOLD responses with increasing dose (linear trend: χ²(1) = 10.80, p = .001; placebo vs. high-dose: β = −0.665, p = .008), with no effect in GT (χ²(2) = 1.978, p = .372; BF₀₁ > 3, indicating evidence for the null). Full results are reported in the Supplementary Methods (Supplementary Fig. 9, panel a).

Two converging observations argue against a regression to the mean account of these findings. The drug effect in ST followed a significant linear dose-response gradient, which is against the odds for a dose-independent statistical artifact. In addition, GT individuals, who showed higher outcome BOLD under placebo, exhibited no reduction in outcome BOLD under drug, and ST individuals, who showed lower outcome BOLD under placebo, exhibited no increase. These results are inconsistent with the symmetric pattern that regression to the mean would predict.

##### Self-reported drug effects

Linear mixed-effects models of three factors derived from Visual Analogue Scales (Energy, Calm, and Focus; a detailed description is provided in Methods and Supplementary Methods) tested Time × Cluster × Phase (drug vs. placebo session) effects for each factor. A significant three-way interaction emerged for Energy only (F(1, 90) = 4.20, p = .043, η²p = .04; Supplementary Fig. 9, panel b). Follow-up analyses within the drug session revealed a significant Time × Cluster interaction (F(1, 30) = 20.47, p < .001, η²p = .41), with Sidak-adjusted post-hoc contrasts (coded as pre minus post-dose, such that positive values indicate a decrease over time) showing a significant decrease in Energy from pre- to post-dose in ST (β = 0.76, SE = 0.14, t(30) = 5.37, p < .001, d = 2.29) and no change in GT (β = −0.03, SE = 0.10, t(30) = −0.29, p = .997, d = −0.09). No Time × Cluster interaction was observed in the placebo session (F(1, 30) = 0.0006, p = .980, η²p < .001), and no significant three-way interactions emerged for Calm (F(1, 90) = 0.59, p = .444) or Focus (F(1, 90) = 0.004, p = .949).

#### Repeated-dose D_2_/D_3_ antagonist and partial agonist cohort

##### Cluster replicability

The repeated-dose D_2_/D_3_ antagonist and partial agonist cohort included 48 healthy participants (26 females, 22 males; mean age = 26.5 years, SD = 8.15) from a randomized, double-blind, placebo-controlled, crossover study^39^. The study employed a between-subjects pharmacological design in which participants received treatment periods with placebo and either amisulpride (a D_2_/D_3_ receptor antagonist) or aripiprazole (a D_2_/D_3_ receptor partial agonist) for 7 days each with a pseudo-randomised counter-balanced order. More details on this cohort and study design are provided in Supplementary Methods.

Discovery cohort ST/GT centroids were projected onto the repeated-dose cohort to obtain predicted cluster labels, and an independent within-sample hierarchical k-means clustering (k=2) was performed to obtain data-driven labels. Agreement between the two methods was assessed via confusion matrix metrics (accuracy, sensitivity, specificity). In the amisulpride arm, overall accuracy was 78.3%, with perfect sensitivity (100%) but lower specificity (50%), and 75.0% replicability based on sensitivity and specificity. In the aripiprazole arm, overall accuracy was 88.0%, with perfect sensitivity (100%) and specificity of 75.0%, and 87.5% replicability. Participants whose labels were predicted and those data-driven were discordant were considered misclassified and excluded from subsequent analyses (amisulpride: N=5; aripiprazole: N=3), leaving a final concordant sample of N=40 (amisulpride: N=18, ST=13, GT=5; aripiprazole: N=22, ST=13, GT=9).

##### Placebo vs. Drug effects on ventral striatal reactivity

Ventral striatum BOLD responses were analyzed using linear mixed-effects models with a random intercept per subject. Within each drug arm, participants completed the MID task under both placebo and active drug conditions. Response values were standardized within each task phase to place anticipation and outcome contrasts on a common scale. All models included age, sex, drug administration order, and baseline BOLD as covariates. A combined model was fitted with treatment condition (placebo, drug), task phase (anticipation, outcome), cluster (ST, GT), and drug arm (amisulpride, aripiprazole) as fixed factors. Because amisulpride and aripiprazole were administered to independent groups with distinct pharmacological mechanisms, per-arm simple effects were estimated from within-phase models retaining drug arm as a factor, consistent with the use of a priori planned comparisons in factorial designs^13,52^.

The four-way Phase × Treatment × Cluster × Drug Arm interaction was not significant (F(1, 136) = 0.34, p = .562, η²p = .002). The three-way Phase × Treatment × Cluster interaction was significant (F(1, 136) = 9.80, p = .002, η²p = .067), indicating that treatment effects on ST and GT differed across task phases.

In the anticipation phase, the Treatment × Cluster interaction was significant (F(1, 37) = 5.69, p = .022, η²p = .133). Amisulpride reduced anticipatory ventral striatum BOLD in ST (β = −0.98, 95% CI [−1.62, −0.35], p = .003, d = 1.07) but not GT (β = −0.19, 95% CI [−1.16, 0.79], p = .703, d = 0.13). Aripiprazole increased anticipatory BOLD in GT (β = 0.86, 95% CI [0.13, 1.58], p = .023, d = 0.81) but had no effect in ST (β = 0.05, 95% CI [−0.56, 0.65], p = .877, d = 0.05). Results are depicted in Figure 3, panel b.

In the outcome phase, the Treatment × Cluster interaction was also significant (F(1, 69) = 8.20, p = .006, η²p = .106). GT showed reduced outcome BOLD under drug relative to placebo across both arms (β = −0.96, 95% CI [−1.67, −0.25], p = .009, d = 0.90), with no change in ST (β = 0.29, 95% CI [−0.24, 0.82], p = .272, d = 0.37). This effect was driven by aripiprazole (GT: β = −1.07, 95% CI [−1.97, −0.17], p = .021, d = 0.82; ST: p = .193), with no significant effects of amisulpride in either cluster (ST: p = .841; GT: p = .214). Results are shown in Figure 3, panel c. Together, these results show that ST and GT respond in qualitatively distinct ways to D_2_/D_3_ antagonism and partial agonism, with amisulpride and aripiprazole producing opposite-sign effects on anticipatory BOLD in the same placebo-defined clusters. This cross-compound divergence further argues against a regression-to-the-mean account, which would predict consistent attenuation toward the cohort mean regardless of pharmacology. Across both pharmacological cohorts, these drug effects held whether non-concordant participants were removed or retained and across the projection and within-sample clustering labeling examined (Supplementary Table 4).

### Clinical cohort

#### Cluster replicability and clinical assessment

Having established that ST and GT phenotypes show dissociable sensitivity to dopaminergic modulation in healthy volunteers under both single- and repeated-dose pharmacological challenge, we next extended this framework to a clinical sample of patients receiving sustained antipsychotic treatment. The clinical cohort consisted of 34 individuals (22 males; mean age = 24.7 years, SD = 5.6) recruited at UNIBA, including 17 with first-episode psychosis (FEP; 11 with Psychotic Disorder NOS, 2 with Brief Psychotic Disorder, 1 with Brief Psychotic Disorder/Mood Disorder NOS, 1 with Paranoid Type Schizophrenia, and 2 with an unspecified diagnosis) and 17 with schizophrenia (SCZ), confirmed by the Structured Clinical Interview for DSM Disorders (SCID). Cluster replicability in this cohort showed good clustering performance (Figure 4, panel a), with 77.1% overall accuracy (sensitivity: 66.7%; specificity: 100%) and 83.3% replicability. While all GT were correctly identified, eight ST were misidentified (mis-ST) and were excluded from subsequent analyses. After exclusion of mis-ST, the final analytic sample included 26 patients (ST: N = 15, with 5 FEP and 10 SCZ; GT: N = 11, with 7 FEP and 4 SCZ).

**Figure 4.**
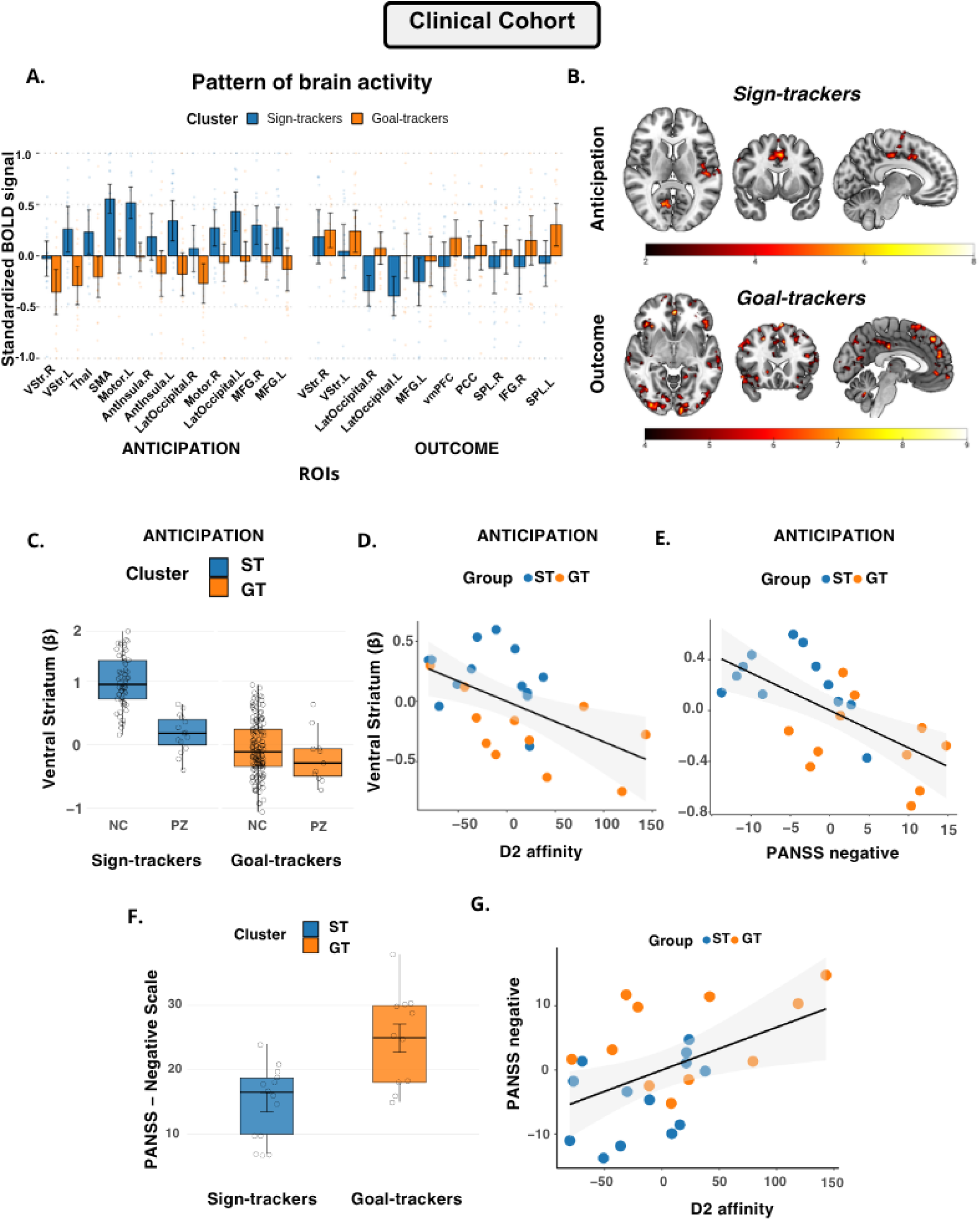
Clinical cohort. **a. Pattern of brain activity.** Concordant cluster assignments (data-driven within-sample k = 2 cross-tabulated with Discovery centroid projection) identified sign-trackers (ST; N = 15) and goal-trackers (GT; N = 11), with the remaining 8 patients excluded as mis-ST. ST showed heightened BOLD activity during reward anticipation but not during outcome, while GT showed greater BOLD activity at reward outcome rather than anticipation. Bars represent 95% CI. **b. Voxel-wise activation maps.** Whole-brain voxel-wise analyses (pTFCE-FWE < 0.05) showed activation in ST during anticipation and in GT during outcome, with no significant activation in the alternate phase. **c. Ventral striatal BOLD during anticipation.** Boxplots show standardized BOLD responses in the ventral striatum (vStr) during reward anticipation, separately for neurotypical controls (NC) from the Replication cohort and patients, within each cluster (ST, GT). **d. D₂ affinity and ventral striatal BOLD.** Scatterplot showing the negative correlation between vStr BOLD during anticipation and the patient-specific D₂ dopaminergic affinity index. **e. Negative symptoms and ventral striatal BOLD.** Scatterplot showing the negative correlation between vStr BOLD during anticipation and negative symptom severity (PANSS Negative Scale). **f. Negative symptoms by cluster.** Boxplots showing PANSS Negative Scale scores in ST and GT patients. **g. D₂ affinity and negative symptoms.** Scatterplot showing the positive correlation between the patient-specific D₂ dopaminergic affinity index and negative symptom severity. *Note.* Scatterplots in panels D, E, and G display residualized data, adjusted for age and sex.

Whole-brain voxel-wise analyses (p_TFCE-FWE_<0.05) showed activation in ST during anticipation and activation in GT during outcome (Figure 4, panel b), with no significant activation in the alternate phase (Supplementary Table 2). When comparing patients to neurotypical controls (NC) derived from the Replication cohort (collected with the same imaging protocol at UNIBA), ST patients showed a lower BOLD signal in the ventral striatum during anticipation, t(71)=7.12, p<0.001, Cohen’s d=2.06 (Figure 4, panel c). In contrast, GT patients did not differ significantly from NC, t(153)=1.02, p=.308, Cohen’s d=0.32. Within patients, ST showed larger ventral striatal BOLD during reward anticipation than GT (t(24) = 2.72, p = .012, d = 1.08), preserving the same direction observed in healthy controls (t(200) = 16.02, p < .001, d = 2.49).

To verify that the ST/GT distinction was not driven by differential medication exposure, we compared chlorpromazine equivalents between groups (data available for 24 of 26 patients) and found no significant difference (p = .50). In this same subset of patients with documented ongoing treatment (N = 24), we calculated a patient-specific D₂ dopaminergic affinity index by integrating individual medication dosages with established receptor binding affinities^53^. Consistent with our hypothesis, this index of D₂ receptor blockade was significantly and negatively correlated with striatal BOLD activity during reward anticipation (r = −0.462, p = .023, Figure 4, panel d), after accounting for age and sex. Affinity for other dopamine receptors yielded no significant associations with striatal BOLD activity in anticipation (D_1_: p = .417; D_3_: p = .105; D_4_: p = .323; D_5_: p = .245).

We next examined whether the same dopaminergic mechanism identified in the healthy pharmacological cohorts left a detectable signature on clinical phenotype expression. Differences in clinical symptoms, assessed through the Positive and Negative Syndrome Scale (PANSS), were tested in association with BOLD signal and D₂ dopaminergic affinity. We found that the BOLD signal in the ventral striatum during anticipation was significantly and specifically associated with negative symptoms (r=−0.579, p=0.002, Figure 4, panel e), while no association was found with either positive symptoms (r=0.03, p=0.901) or general psychopathology items (r=−0.15, p=0.486). Consistent with these findings, GT individuals demonstrated significantly higher negative symptom severity than ST individuals (Wilcoxon W=144, p<0.001, Figure 4, panel f). Furthermore, a significant positive association was found between the D₂ dopaminergic affinity index of the administered therapy and negative symptom severity (r=0.471, p=0.020, Figure 4, panel g). No other significant associations were detected between striatal BOLD activity in anticipation and dopaminergic affinity (D_1_: p=0.336; D_3_: p=0.244; D_4_: p=0.140; D_5_: p=0.320). Together, these results suggest that patients administered antipsychotic treatments characterized by stronger D₂ blockade also show a GT imaging phenotype and have a higher negative symptom burden. These associations and the negative-symptom difference between phenotypes held whether non-concordant patients were removed or all were retained (Supplementary Table 4).

## DISCUSSION

We translated the well-characterized rodent ST/GT framework to humans using neuroimaging to investigate individual differences in reward processing and the effects of antipsychotics with high affinity for D₂ receptor modulation. While previous human studies have explored ST/GT analogs with different approaches^15,20^, our study provides a translationally relevant validation grounded in the functional dissociation between reward anticipation and outcome phases, which maps onto the core mechanistic distinction between ST and GT neural profiles, across five independent cohorts. Our results highlight four main findings. First, the ST/GT model is highly replicable in humans even without new learning, as BOLD signal clustering consistently delineated ST-like and GT-like profiles across all cohorts. Second, ST attribute greater incentive salience to reward-anticipation cues, showing enhanced ventral striatal and whole-network responses during anticipation, whereas GT show preferential activation during outcome processing. Trial-by-trial analyses further confirmed that this elevated anticipatory response is sustained across trials and does not require active learning to be maintained. Third, full D₂/D₃ antagonism selectively attenuated anticipatory ventral striatal BOLD in ST, with a corresponding decrease in self-reported Energy, across both single-dose (risperidone, haloperidol) and seven-day (amisulpride) administration, while a partial agonist (aripiprazole) produced a qualitatively distinct, phenotype-specific pattern. Finally, in patients with psychosis, those treated with strong dopaminergic antagonists showed the most blunted ventral striatal anticipation signals and the highest negative-symptom burden. Together, these findings demonstrate a translational pathway from preclinical models to healthy humans and, ultimately, to clinical populations, establishing a neuroimaging biomarker of incentive salience and offering a phenotype-driven strategy for tailoring D₂-targeted interventions.

In the Replication cohort, human ST showed superior executive flexibility on the WCST, with fewer perseverative and non-perseverative errors and more efficient contingency switching, partially convergent with the enhanced set-shifting reported in ST rodents^28^ but inconsistent with their deficits in response inhibition. These mixed correspondences likely reflect divergent task demands across species, with human assays emphasizing explicit rule abstraction and rodent assays relying on implicit Pavlovian-instrumental transfer^11^.

Beyond replicability, the pharmacological results provide a cross-species test of convergent validity for the human imaging ST/GT phenotypes, asking whether they respond to dopaminergic manipulation in the same way as the rodent behavioral phenotypes from which the framework was derived. That manipulating dopaminergic signaling in humans alters BOLD responses during reward anticipation is by now well established, with evidence from D₂-receptor antagonists (e.g., risperidone^36^, amisulpride^54^, haloperidol^41^, olanzapine^55^, lurasidone^56^, sulpiride^57^), indirect dopamine agonists (dextroamphetamine^58^, amphetamine^59^), and the dopaminergic precursor L-DOPA^41^. What is novel here is that, in humans, ST exhibit a selective response and dose-dependent attenuation of BOLD activity after challenge with antipsychotics that have high affinity for D₂ receptor antagonism, accompanied by a parallel decrease in self-reported Energy. The effect on self-reports is especially noteworthy, given the potential of fMRI-mediated ST-GT discrimination to predict treatment response and side effects of dopaminergic agents. This specific dopaminergic sensitivity in ST mirrors findings from rodent work showing that cue-driven incentive salience in ST rats depends critically on dopamine transmission^29,30^. By contrast, GT rats acquire cue–reward associations with comparatively less dopaminergic involvement. Together, these results underscore the central role of dopamine in modulating incentive salience^60,61^ and demonstrate that ST individuals, who attribute greater motivational value to cues than GT, are particularly sensitive to high-affinity D₂ receptor antagonists at both the neural and subjective levels.

Building on the single-dose results, we extended the pharmacological logic to repeated-dose administration of two D_2_/D_3_ agents with mechanistically distinct profiles, asking whether the ST-specific sensitivity generalized beyond acute exposure and whether it depended on full antagonism. Repeated-dose amisulpride, a selective D_2_/D_3_ antagonist, replicated the single-dose pattern: anticipatory ventral striatal BOLD was selectively reduced in ST but not in GT. This convergence across acute (risperidone, haloperidol) and seven-day (amisulpride) full antagonism is notable because the original group-level analysis of the same dataset reported no significant effect of amisulpride on reward anticipation in the pre-specified striatal ROI^39^; the ST/GT phenotyping recovered an effect that was masked when participants were considered as a unitary group. This finding extends prior demonstrations that compounds with D_2_ receptor antagonism reduce dopamine-dependent reward signals in healthy humans^36,41,55^ by suggesting that the underlying sensitivity is not uniformly distributed but rather bimodal. By contrast, repeated-dose aripiprazole, a D_2_/D_3_ partial agonist, produced a qualitatively different pattern, with increased anticipatory ventral striatal BOLD in GT and reduced outcome-phase BOLD in GT, while ST responses were largely unchanged; because this dissociation derives from a single cohort and contrasts with the consistent ST-specific attenuation observed under full antagonism, it should be interpreted as preliminary and warrants replication. Together, these results suggest that the null group-level findings reported by Osugo et al.^39^ may not reflect an absence of pharmacological action on reward processing, but rather phenotype-specific effects that become apparent only once ST and GT individuals are separated.

Our findings inform the aberrant salience hypothesis of psychosis, which proposes that dysregulated dopamine signaling drives inappropriate significance attribution and that antipsychotics ameliorate symptoms by normalizing these signals^62^. In our clinical cohort, hierarchical clustering again identified ST and GT profiles despite the administration of antipsychotic medications. However, even ST patients showed markedly blunted ventral-striatal responses during reward anticipation compared with healthy ST, suggesting that prolonged antipsychotic treatments may mask intrinsic cue-driven activation. In contrast, GT patients did not exhibit the same blunted activation profile compared to healthy GT individuals. This finding is noteworthy because it suggests that the classic “blunted” striatal response often described in schizophrenia^40^ is not a universal characteristic of the disorder, but a variable feature that our ST/GT framework may help explain. This perspective reinforces the argument that we should expect different treatment responses based on phenotype. Furthermore, while conventional antipsychotics primarily target positive symptoms, the fact that our ST/GT phenotyping was linked to an effect on negative symptom burden, a dimension often resistant to treatment, opens the possibility of a phenotype-driven strategy to uncover a neurobiological mechanism of an important unmet need in psychiatry.

When dealing with blunted striatal activity, disentangling trait-like salience attribution from drug-induced effects is challenging. On the one hand, patients classified as GT by their BOLD signature may truly reflect goal-tracking phenotypes; on the other, long-term antipsychotic exposure itself remodels striatal function. Benjamin et al.^63^ reported that chronic D₂ antagonism drives extensive transcriptional changes in the human caudate, including alterations in *DRD2*-related pathways. Our findings position this concern within an empirical timescale: with phenotypes defined under placebo in both pharmacological cohorts, the ST-specific anticipatory reduction was detectable both after single-dose challenge with risperidone and haloperidol and after seven days of repeated-dose amisulpride, indicating that the phenotype-specific sensitivity to D_2_/D_3_ antagonism is not restricted to acute receptor occupancy and is already present at sub-chronic exposure windows. In patients receiving sustained antipsychotic treatment for months to years, this acute and sub-chronic suppression is plausibly compounded by slower, gene-expression-mediated remodeling of the reward circuitry, producing the more pronounced and stable blunting characteristic of the clinical state. The blunted anticipatory BOLD response observed in our GT-like patients, in conjunction with their higher negative symptom burden and a positive correlation between D₂ affinity and negative symptoms, is consistent with the interpretation that the observed neural and behavioral deficits may partly reflect the cumulative consequences of long-term treatment with high-affinity D₂ antagonists.

Across cohorts, the data were consistent with two phenotypes rather than three. The sign- and goal-tracking phenotypes were recovered whether participants were grouped by clustering or by a fixed criterion on the first principal component, and we interpret the participants situated between them not as a distinct third group but as the less prototypical members of the two distributions, which are hard to distinguish in terms of the underlying group membership. This is consistent with the foundational operationalization of sign- and goal-tracking, in which a fixed and admittedly arbitrary criterion on a tracking index defines an intermediate group that is commonly set aside from the sign-versus goal-tracker comparison^30,64^, an approach carried into its human translation^13,15^. That criterion is acknowledged to be arbitrary and to differ across laboratories^64^, and the fraction of animals falling in the intermediate region varies across populations of the same strain^65^, so its size reflects where the boundary is drawn rather than the presence of a third mode. Across the Discovery, Pharmacological, and Clinical cohorts, the principal effects were broadly similar whether the phenotypes were defined by the three-group or the two-group solution, and whether non-concordant participants were removed or retained, and the Discovery neuropsychological dissociation was preserved across a range of intermediate widths. The intermediate region is also where measurement precision and detection power are lowest, so the present data constrain the structure that can be resolved rather than settle whether a distinct intermediate phenotype exists at a finer grain than these measurements allow. We therefore regard the intermediate participants as the less prototypical members of the two distributions rather than as a separate phenotype, an interpretation of what the present data can resolve rather than a claim that no such phenotype exists.

Despite the robust replicability of our clustering approach and the clear correspondence between our behavioral and neural findings and those from animal models, several limitations warrant consideration. First, the absence of subjective reports on whether participants were aware of the treatment condition during each session, although the observed ST-specific reduction in Energy suggests that drug effects were behaviorally meaningful despite this uncertainty. Second, the lack of clinical scales to supplement the self-reported data in the Pharmacological cohorts is a limitation. Third, our modest patient sample, which included both first-episode and chronic schizophrenia cases, limits our ability to definitively disentangle trait-like salience attribution from treatment effects. The challenge of separating intrinsic phenotypes from drug-induced brain adaptations is particularly relevant here, as long-term antipsychotic exposure can remodel striatal function. This leaves open the compelling question of whether patients exhibiting a GT-like activation profile are those receiving ongoing therapy, rather than reflecting a distinct underlying phenotype. Fourth, the single-dose cohort comprised only healthy males, which limits the generalizability of the single-dose ST-specific D_2_ antagonism effect, given known sex differences in pharmacokinetics and dopaminergic signaling^66,67^; the repeated-dose cohort, which included both sexes, partially addresses this concern. Fifth, risperidone and haloperidol collapsed into a single drug condition due to sample size constraints, which may introduce heterogeneity given differences in receptor binding profiles^68^. The dose-response analysis conducted separately in the risperidone subsample is consistent with the primary analysis and partially mitigates this concern.

In summary, this study identifies robust ST and GT imaging phenotypes in humans, mirroring rodent models of reward processing. ST exhibit heightened anticipatory striatal activity and dopamine-sensitive cue reactivity, whereas GT show preferential engagement during reward outcome processing. These profiles were consistently detected across tasks/cohorts, drug conditions, and clinical status, linking them to both subjective drug response and symptom severity in psychosis. By translating preclinical models into a replicable human framework, our findings offer a neuroimaging biomarker of incentive salience and a promising lead into fMRI-based clinical predictions.

## MATERIALS AND METHODS

### Participants

The study included fMRI data from 1,251 individuals (described in the Supplementary Methods and Supplementary Table 1), comprising a Discovery cohort of 890 healthy participants from IMAGEN^42^ (454 females, 436 males; mean age=22.1 years, SD=.7 years), a multi-site European study, an independent Replication cohort of 245 healthy individuals collected at the University of Bari Aldo Moro (UNIBA; 141 females, 104 males; mean age=25.7 years, SD=5.7), a Pharmacological Challenge cohort including two datasets: *Single-dose* D_2_ antagonists, of 34 healthy males (mean age=26.9 years, SD=6.8); *Repeated-dose* D_2_/D_3_ antagonist and partial agonist, of 48 healthy adults (26 females, 22 males; mean age=26.5 years, SD=8.15), and a Clinical cohort of 34 individuals collected at UNIBA (12 females, 22 males; mean age=24.7 years, SD=5.6), including 17 with FEP and 17 with SCZ confirmed by the SCID. Inclusion criteria were the same as those for the healthy subjects enrolled in the independent Replication cohort and are fully detailed in Supplementary Methods.

While the Discovery and Replication cohorts included only a single time point per participant, the *single-dose* cohort participated in a double-blind, placebo-controlled, three-period, crossover study. Specifically, for the *single-dose* cohort, data from two studies were considered here^36,37^, in which participants were pseudo-randomly assigned to receive either risperidone (0.5 mg or 2 mg) and placebo (N=17) or haloperidol (3 mg), and placebo (N=17), undergoing three MRI sessions at one-week intervals (a detailed description of the sample and the design is provided in the Supplementary Methods). The *repeated-dose* D_2_/D_3_ antagonist and partial agonist cohort participated in a single-center, randomized, double-blind, placebo-controlled, crossover study. Two independent groups of healthy volunteers received either amisulpride and placebo (arm 1) or aripiprazole and placebo (arm 2) for 7 days each. The study employed a between-subjects pharmacological design in which participants received either amisulpride (a D_2_/D_3_ receptor antagonist) or aripiprazole (a D_2_/D_3_ receptor partial agonist).

### Experimental design

As illustrated in Figure 1, panel a, the study flowchart comprises neuroimaging acquisition, neuropsychological assessment, self-report, and clinical assessment.

All five cohorts employed a variant of the MID task (Figure 1, panel b), a widely used paradigm for investigating the neural correlates of reward anticipation and outcome processing^27,69^. A common feature across cohorts is that the target duration or response window is calibrated, either online or from training-phase performance, to maintain a comparable hit rate across participants (Supplementary Methods provide cohort-specific parameters), thereby preventing BOLD differences across phenotypes from being confounded by phenotype-related differences in task accuracy. Details on task design, fMRI data acquisition, and statistical processing are reported for each cohort in the Supplementary Methods.

In the Discovery cohort, the neuropsychological assessment included evaluations of personality and cognitive function. To assess lower order trait dimensions more specifically related to disinhibitory psychopathology, a total of five scores from 35 self-reported TCI-R^44,45^ items evaluating only the Novelty Seeking, as the tendency to respond impulsively to novel stimuli, were included in the analyses, i.e., the overall Novelty Seeking score and four scores for (Exploratory excitability, Impulsiveness, Extravagance, and Disorderliness) corresponding to its subcomponents according to the Cloninger’s model^70^. To evaluate the potential risk factors for addictive behaviors and comorbid psychopathology development, four scores, each for one personality trait, from the SURPS^46^ were computed for analysis purposes: hopelessness (as the tendency to develop bleak expectations about oneself and the future), anxiety sensitivity (as the fear of anxiety-related physical sensations), impulsivity (as lack of premeditation and difficulties with response inhibition), and sensation seeking (as the need for intense and novel experiences).

To evaluate decision-making, we examined six CGT scores: Delay Aversion, Deliberation Time, Overall Proportion Bet, Quality of Decision Making, Risk Adjustment, and Risk Taking. Finally, to determine passive avoidance learning, nine outcome measures from the PALP^47^ were included in the analysis: the number of commission errors, the number of omission errors, and reaction times for each of the following reward contingencies: PP=Punishment-Punishment; RP=Reward-Punishment; RR=Reward-Reward.

In the Replication cohort, we assessed personality using TCI^45^, obtaining seven scores, namely Novelty Seeking, Harm Avoidance, Reward Dependence, Persistence, Self-Directedness, Cooperativeness, and Self-Transcendence. To replicate some of the cognitive processes elicited by the PALP task, e.g., set shifting, inhibitory control, and contingency learning, we estimated four cognitive scores from the WCST^50^, recorded as the number of completed categories, number of trials, and number of perseverative and non-perseverative errors. Finally, the following scores from the CPT-AX^49^ evaluating sustained attention and cognitive control were computed and included in the analysis: the ratio of the target stimuli detected (Hits), the ratio of the false alarms, i.e., the inappropriate responses given to a distractor appearing after the letter “A” (FA)^17^ and “B” (FAC), a sensitivity index calculated using the proportion of total false alarms (AP S), the proportion of AY errors (AY), individual’s ability to discriminate signal from noise (DP S), and sensitivity of target detection in relation to the nature of the preceding stimulus (DPC S).

In the Pharmacological Challenge cohort (*Single-dose*), participants completed a battery of 16 visual analog scales (VAS) designed to assess their momentary cognitive and affective state. Each scale consisted of a 100-mm horizontal line anchored by opposing descriptors, with participants indicating their current experience by marking a point along the line. Scores were recorded in millimeters from the left (0 mm) to the right (100 mm), such that higher values reflected greater endorsement of the right-side descriptor. The 16 VAS items included the following pairs: Alert–Drowsy, Energetic–Tired, Strong–Feeble, Motivated–Unmotivated, Determined–Aimless, Focused–Distracted, Clear-headed–Mentally foggy, Able to concentrate–Unable to concentrate, Calm–Stressed, Relaxed–Anxious, At ease–On edge, Confident–Unsure, In control–Overwhelmed, Sociable–Withdrawn, Cooperative–Resistant, and Friendly–Irritable. Ratings were collected at two time points: immediately before drug or placebo administration (pre-dose) and again 90 minutes later (post-dose), approximately 30 minutes before the MRI session.

In the Clinical cohort, treatment information was available for 24 out of 26 in the final sample on ST/GT. For these patients, the chlorpromazine equivalent was computed using the method described by Gardner et al.^71^. In this sample, clinical symptoms were assessed during a clinical evaluation preceding the MRI session using the PANSS.

Further details on the neuropsychological assessment in the Discovery and Replication cohorts, as well as the self-reported assessment in the S*ingle-dose* Pharmacological challenge cohort and the clinical assessment in the Clinical cohort, are provided in the Supplementary Methods.

### Statistical Analyses

We developed a multi-step pipeline that integrates neuroimaging, machine-learning analyses, and pharmacological challenges to identify imaging phenotypes of reward-related brain activity and to validate them across independent cohorts.

First, meta-ROIs were defined from a published meta-analysis of 81 studies that did not include the current datasets^6^ (Figure 1, panel b). These ROIs captured key regions implicated in reward processing, such as the ventral striatum during anticipation and the medial prefrontal cortex during outcome^72^ (Supplementary Fig. 1). Using Marsbar ^73^, the mean BOLD signal was extracted for each individual from twelve ROIs during anticipation and ten ROIs during outcome. This procedure was applied consistently across all cohorts.

As depicted schematically in Figure 1, panel d, clustering was first performed on the Discovery cohort to examine whether participants could be grouped based on reward-related activity. Hierarchical k-means clustering was applied to the whole dataset, using Euclidean distance and Ward’s linkage as implemented in the factoextra R package^74^. Outliers were identified with Rosner’s generalized extreme Studentized deviate test^75^ and iteratively removed. All features were scaled to the range -1 to 1, and the regression residuals accounted for site effects. The optimal number of clusters (*k* = 3) was determined using two complementary approaches: the *NbClust* R package and a hypothesis-driven approach. We nonetheless set *k* = 3 on hypothesis-driven grounds, following prior work on the ST/GT model^13,76^, to recover the ST, GT, and INT phenotypes. Resolving the intermediate group as a distinct cluster was intended to define the ST and GT centroids on their more prototypical members, so that the centroids projected onto the external cohorts were not displaced toward the center of the distribution by transitional cases. The separation of the two phenotypes was further supported by Ashman’s D and the Mahalanobis distance between the ST and GT centroids, as visualized in principal component space (see Supplementary Methods and Supplementary Figs. 2-3). To validate this solution, we implemented a leave-site-out framework in which the Discovery dataset was split into a training set comprising 7 sites and a test set comprising 1 site. Clustering was performed on the training set, and predicted memberships were assigned to the test set using only the ST/GT centroids. Validation was assessed using confusion matrices^77^, comparing predicted cluster assignments with those obtained through independent clustering in the testing set.

The Discovery model was then applied to the Replication, Pharmacological Challenge, and Clinical cohorts (Figure 1, panel d). In each case, data were scaled to the same range, clustering was repeated, and predicted assignments were compared with independent clustering results via confusion matrices. Sensitivity, specificity, and balanced accuracy were computed, using balanced accuracy as the primary measure of replicability. Balanced accuracy values were interpreted against the conventional thresholds for clinical prediction (≥0.70 acceptable, ≥0.80 good^78^) and benchmarked against published fMRI-derived psychiatric classifiers^79^. Rather than adopting a single fixed threshold, classification performance was evaluated across multiple independent cohorts, reasoning that consistent above-chance accuracy replicated across datasets provides stronger evidence for phenotype validity than any single cutoff applied to one sample.

Next, we examined differences between ST and GT in personality and cognitive functioning using Wilcoxon rank-sum tests in the Discovery and Replication cohorts. In the Replication cohort, the WCST was analyzed using a one-tailed Wilcoxon test to test the hypothesis that ST would outperform GT and to reproduce the effect direction observed in PALP results. All results were corrected for multiple comparisons with a threshold of p_FDR_ < 0.05.

In the Pharmacological Challenge cohorts, analyses addressed both neuroimaging measures and self-reported drug effects. Differences in self-reported Energy, Calm, and Focus, derived from confirmatory factor analysis of 16 visual analog scale items, were modeled with linear mixed-effects models that included fixed effects for Time, Cluster, and Phase (Placebo versus Drug), as well as their interactions, while adjusting for drug type (risperidone versus haloperidol) and age. ROI-level BOLD signals were averaged within each phase and analyzed with mixed-effects models including Phase, Condition, and Cluster as fixed effects, with drug type and age as covariates.

Dose-response analyses of risperidone were then conducted with linear and quadratic mixed-effects models, and model selection was based on likelihood ratio tests. Significance of post hoc comparisons was set at p_FDR_ < 0.05. Additional robustness checks included Levene’s test for variance homogeneity, which confirmed equivalent BOLD variance between ST and GT under placebo (F(1,32)=0.54, p=0.468), and Bayesian analyses (BF₀₁) to quantify evidence for the null hypothesis in the absence of significant effects. Full details of these robustness checks and the dose-response analyses are reported in the Supplementary Methods.

In the Repeated-dose D_2_/D_3_ antagonist and partial agonist sub-cohort, the linear mixed-effects models reported in the Results were estimated with the following analytical choices. Response values were standardized within each task phase prior to modeling, in order to place the anticipation and outcome contrasts on a common scale and allow direct comparison of effect magnitudes across phases; this within-phase standardization is a linear transformation and does not alter within-phase test statistics. Four covariates were included in all models: age, sex, drug administration order, and baseline BOLD response. Drug administration order was retained because, in this dataset, the relatively short washout interval between sessions could, in principle, introduce order-dependent variability in placebo or drug responses; baseline BOLD response was included to account for between-subject differences in pre-treatment ventral striatal reactivity. Statistical significance was assessed using Type III ANOVA with Satterthwaite degrees of freedom; effect sizes are reported as partial η² for omnibus interactions and Cohen’s d for cell-wise contrasts. In addition to the combined and per-phase models reported in the main Results, eight separate linear mixed-effects models were fitted (one per drug arm × phase × phenotype combination) to test within-cell drug effects, with placebo–drug contrasts FDR-corrected (Benjamini–Hochberg) within phase and across all eight tests; per-cell results are provided in the Supplementary Methods.

In the Clinical cohort, we integrated antipsychotic receptor binding profiles with clinical dosing parameters to compute dose-normalized receptor affinity ratios. Equilibrium dissociation constants (Ki values) for dopamine receptors D1–D5 were obtained from the dataset published by McCutcheon et al.^53^ via their public GitHub repository, while clinical data comprised patient-level medication regimens, individual dosages, and chlorpromazine-equivalent doses (CPZ). Dose-to-affinity ratios were calculated as CPZ equivalents divided by the respective Ki values (nM) for each receptor subtype, yielding dimensionless indices of receptor occupancy potential at therapeutic doses. These indices quantitatively measure the relative blockade capacity for each receptor subtype under clinically relevant dosing conditions and were generated using a custom R pipeline. The final analytic sample included 24 patients, after excluding individuals with missing data or misidentification in clustering. Group differences in ventral striatum BOLD signal between patients and healthy controls from the UNIBA cohort were tested with independent-sample t-tests. In contrast, ST–GT differences in positive and negative symptoms were examined using Wilcoxon rank-sum tests (N=25, with PANSS data unavailable for one patient). Correlational analyses further assessed associations between receptor affinity indices, ventral striatum activity, and clinical symptom dimensions.

All analyses were performed in RStudio (2023.09.1+494), and additional details on preprocessing and statistical procedures are available in the Supplementary Methods.

## Supporting information

Supplementary Informations

## Data availability

Data from IMAGEN are available from a dedicated database at https://imagen-project.org. Data from UNIBA cannot be shared at the individual level because of ethical restrictions based on the protocol approved by the relative institutional ethics committees to protect the privacy of the participants. Data from King’s College London corresponding to the *Single-dose* Pharmacological Challenge cohort using single-dose D_2_ antagonists can be requested to the corresponding author Mithul A. Mehta. De-identified individual participant data corresponding to the *Repeated-dose* Pharmacological Challenge cohort are available for research purposes from Martin Osugo from the publication date, subject to a data-sharing agreement, except for data from a minority of subjects who did not consent to de-identified data being used to support future research.

## Code availability

Activation maps at the group level for the Discovery, Replication, and Clinical cohorts are available along with meta-ROIs and the R code used to perform cluster analysis at the repository https://github.com/nsambuco/MID_ST-GT.

## Acknowledgments

A.L. is a Ph.D. student in the National Ph.D. in Artificial Intelligence, XXXVIII cycle, Health, and Life Sciences course, organized by Università Campus Bio-Medico di Roma. The authors would like to thank Yu Chen (Department of Psychiatry, Yale University School of Medicine, New Haven, CT 06520, USA) for sharing the meta-analytic maps used in the current study. The authors are grateful to Prof. Daniel R. Weinberger and Prof. Juergen Dukart for providing scientific insight contributing to this work.

## Author contributions

Conceptualization: NS, AL, GP

Methodology: NS, AL, PCTH, PS, MS, MAM, GP

Investigation: PCTH, AB, GB, LG, DG, FM, AAM, AR, TB, GJB, ALWB, RB, SD, HF, HG, PG, AG, AH, JLM, MLPM, EA, FN, DPO, LP, MNS, NH, NV, HW, RW, the Apulian Network on Risk for Psychosis, OH, MO, MAM, GP

Visualization: NS, AL

Funding acquisition: LAA, AB, GB, GL, AR, OH, MO, MAM, GP

Project administration: NS, AL, PCTH, OH, MO, MAM, GP

Supervision: MAM, GP

Writing – original draft: NS, AL, MAM, GP

Writing – review & editing: PCTH, LAA, AB, GB, GL, LG, DG, PH, GML, FM, AMM, MO, RP, AR, PS, TB, GJB, ALWB, RB, SD, HF, HG, PG, AG, AH, JLM, MLPM, EA, FN, DPO, LP, MNS, NH, NV, HW, RW, and the Apulian Network on Risk for Psychosis.

