## Supplementary Informations for "Reproducible Human Reward Imaging Phenotypes Exhibit Differential Sensitivity to Dopamine D2 Receptor Antagonism"

### SUPPLEMENTARY METHODS

#### MATERIALS AND METHODS

##### Participants

###### Discovery cohort

The IMAGEN study[1] is a multisite, multinational longitudinal project that was carried out in eight European sites, namely Berlin, Dresden, Dublin, Hamburg, London, Mannheim, Nottingham, and Paris. The study involved adolescents recruited from high schools, and to obtain a diverse sample in terms of socio-economic status, emotional and cognitive development, private, state-funded, and special units were equally targeted. All participants underwent four types of assessment for different data domains (biological samples, brain imaging, clinical characteristics, and functioning data). These assessments were conducted at baseline (14 years of age) and followed up at 16, 19, and 22 years (i.e., follow-up 1, follow-up 2, and follow-up 3, respectively). Exclusion criteria included: the presence of overt neurological conditions such as epilepsy, brain tumors, bacterial infections of the CNS, muscular or myotonic dystrophy; cerebral trauma with loss of consciousness of more than 30 min; developmental issues such as major neurodevelopmental disorders, nutrition and metabolic diseases, uncorrectable visual or auditory deficits, IQ < 70; treatment for schizophrenia or bipolar disorder; presence of medical condition such as type 1 diabetes, systemic rheumatologic disorders, malignant tumors requiring chemotherapy, congenital heart defects or cardiac surgery, aneurysms; pre/perinatal issues such as maternal diabetes during pregnancy, excessive alcohol use of the mother during pregnancy, premature birth < 35 weeks and/or detached placental, hyperbilirubinemia requiring transfusion; MRI contraindication such as the presence of metal or electronic implants and severe claustrophobia. A detailed description of recruitment and research procedures has been previously reported[1].

Clinical and behavioral assessments were performed using Psytools software (Delosis Ltd, London, UK) via its Internet-based platform. The battery of questionnaires and cognitive tasks was self-administered both at participants' homes and at neuroimaging facilities.

For the current study, the discovery sample was composed of a large sample of 1004 healthy young adults within Follow-up 3 (FU3) with Monetary Incentive Delay (MID)[2] task data available. Data from 114 participants were excluded because of poor behavioral performance at the MID task (accuracy below chance). No additional participants were excluded after the MRI quality check[3], e.g., if the mean root-mean-square framewise displacement exceeded 0.5mm (see below in the fMRI Processing section). The final discovery sample included a total of 890 participants (age: M=22.1, SD=.7 years; 454 F 436 M). Demographic details grouped by sites are described in Supplementary Table 1. Participants and their parents provided informed consent. The IMAGEN study protocol was approved by the KCL (King's College London) College Research Ethics Committee CREC/06/07-71 and by local ethics research committees at each site. Parents and adolescents gave written consent and verbal assent, respectively.

###### Replication cohort

We recruited an independent sample of 252 healthy adults who underwent a functional magnetic resonance imaging (fMRI) scan, performing a modified version of the MID task[4] and a neuropsychological and cognitive assessment. One subject was removed at his/her request. After removing six subjects for MRI data quality (e.g., excessive motion) and poor behavioral performance at the MID task (accuracy below chance), the final replication sample included 245 participants (age: M=25.7, SD=5.8; 141 F, 104 M). Participants signed an informed consent form complying with the Declaration of Helsinki after fully explaining all procedures approved by the local ethics committee. All subjects were Caucasians from the region of Apulia, Italy. Inclusion criteria were the absence of any lifetime psychiatric disorder, as evaluated with the Structured Clinical Interview for Diagnostic and Statistical Manual of Mental Disorders IV, NP (Non patient version), of any significant neurological or medical condition revealed by clinical and magnetic resonance imaging evaluation, of history of head trauma with loss of consciousness, and of pharmacological treatment or drug abuse in the past year.

**Pharmacological challenge cohorts**

***Single-dose***

Forty-two healthy right-handed males were pseudo-randomly assigned to one of two parallel groups in a double-blind, placebo-controlled, fully counterbalanced three-period crossover study design. All subjects were scanned three times, with 7 days separating each scan. Scanning was conducted at the same time of day per visit, and participants performed a modified version of the MID task[5]. During each visit, each volunteer received a single capsule given orally with water. In one group the capsule contained either a single dose of risperidone 0.5 mg or risperidone 2 mg or placebo (herein referred to as group RIS-H/L); while in the other group participants received either a single oral dose of olanzapine (7.5 mg) or haloperidol (3 mg) or placebo (herein referred to as OLAN/HAL). Within-group treatment order was randomized using a Williams square design. Two subjects were discounted from the HAL/OLAN group due to DESPOT protocol unavailability, while two additional subjects were removed due to structural or functional image artifacts, leaving 17 subjects in this group (age: M=28.8, SD=6.3). In the RIS-H/L group, four subjects were removed for poor behavioral performance at the MID task (accuracy below chance), leaving a final sample of 17 subjects (age: M=27.6, SD=6.9 years). The study was approved by the London (Brent) Human Research Ethics Committee (REC reference: 13/LO/ 1183). Inclusion criteria required normal ECG, standard laboratory blood screens and urinalysis, and alcohol consumption within the recommended guidelines at the time of the study (<21 units per week). Exclusion criteria included a history of neurological of psychiatric illness, physical illness, and positive drugs of abuse or alcohol breath test on the screening or study days. Three volunteers from each group reported smoking one cigarette per day. However, smoking was not permitted on the study days, so it is unlikely the potential acute effects previously reported would be a factor here[6]. A total of 34 subjects (age range 19–42 years, mean ± SD=26.9±6.8) were included in the final pharmacological challenge cohort.

***Repeated-dose***

Healthy volunteers aged 18–65 years were recruited for a single-centre, randomised, double-blind, placebo-controlled, crossover study[7]. Two independent groups of healthy volunteers received either amisulpride and placebo (arm 1) or aripiprazole and placebo (arm 2) for 7 days each. Within each arm, the order of administration was randomised and counter-balanced to ensure approximately equal numbers received active drug and placebo first.

Seventy-six healthy volunteers with no history of neuropsychiatric disorder were recruited and fifty completed the experiments. There were no significant differences between the amisulpride (n = 25) and aripiprazole (n = 25) samples on any demographic variables or on the proportion receiving placebo first.

Amisulpride doses were titrated up to 400 mg/day (day 1: 200 mg, day 2: 300 mg, days 3-7: 400 mg). Aripiprazole doses were titrated up to 10 mg/day (day 1: 5 mg, day 2: 5 mg, days 3-7: 10 mg). Volunteers were evaluated at a screening appointment prior to randomisation. After the screening visit, eligible subjects were randomised to treatment order (amisulpride or placebo first in arm 1, aripiprazole or placebo first in arm 2). These subjects subsequently returned for the baseline assessment, following which the first dose of study medication was administered at the research facility, and the remaining 6 days of medications were dispensed to be taken at home. After completing the first treatment period, subjects returned for outcome and safety assessment after seven days, before entering a washout period of at least five half-lives of the drug and its active metabolites (minimum 10 days for amisulpride, minimum 28 days for aripiprazole). After the washout period, subjects returned to the research facility and commenced the other treatment condition. Compliance was assessed at the end of each treatment week with pill counts and serum drug levels. Only subjects with detectable amisulpride/aripiprazole levels following the active treatment week were included in the analysis sample (N=50, F 28, M 22, age: M = 26.6, SD = 8.15). This study was approved by the London – West London and GTAC NHS Research Ethics Committee (Ethics Committee Reference Number: 18/ LO/1044). All subjects provided written, informed consent prior to participation. Exclusion criteria were; history of psychiatric illness (including alcohol/substance dependence or abuse, other than caffeine/nicotine) as determined by self-report and the Mini-International Neuropsychiatric Interview; current use of any illicit substances as determined by urine drug of abuse testing and self-report; pregnancy as determined by urine pregnancy testing and self-report; self-report of a first degree relative with a psychotic disorder, current or significant previous use of psychotropic or dopamine modulating drugs, breastfeeding, or participation in a study of unlicensed medicines within the previous 30 days; self-report or clinical findings of significant CNS disorder (e.g. significant head trauma, epilepsy, etc.), significant medical disorder, contraindications to dopamine antagonists/partial agonists or MRI scanning; or clinically relevant abnormal findings at the screening assessment, as determined by the principal investigator. In the current study, the final sample includes forty-eight subjects (F 26, M 22; age: M=26.5, SD=8.15).

**Clinical cohort**

We also recruited 34 individuals, comprising 17 with first-episode psychosis (FEP) and 17 with schizophrenia (SCZ) in the region of Apulia, Italy (12 F 22 M; age: M=24.7, SD=5.6). Recruitment procedures were conducted in accordance with The Code of Ethics of the World Medical Association (Declaration of Helsinki), and approval was given by the local ethics committee ("Comitato Etico Indipendente Locale - Azienda Ospedaliero-Universitaria Consorziale Policlinico di Bari"). Diagnosis of SCZ was made using the Structured Clinical Interview for the DSM-5, Axis 1 disorders by board-certified psychiatrists. Exclusion criteria were: a significant history of drug or alcohol abuse; active drug abuse in the previous year; history of head trauma with loss of consciousness; any other critical medical condition.

**Experimental design**

All five cohorts completed an fMRI scan, performing a variation of the MID task, a paradigm extensively used to investigate the neural correlates of reward anticipation and outcome processing[8]. Both the Discovery and Replication cohorts included a neuropsychological assessment. In the Pharmacological Challenge cohort, participants rated their momentary cognitive-affective state using 16 visual analog scales anchored by opposing descriptors (e.g., Alert–Drowsy, Focused–Distracted, Calm–Stressed). Ratings were collected 90 minutes before and after drug or placebo administration, with higher scores indicating greater endorsement of the right-end descriptor on each scale. Finally, the Clinical cohort employed the standardized clinical interview, i.e., Positive and Negative Syndrome scale (PANSS), to assess symptom severity.

**Neuroimaging task**

The MID task has been extensively used to elicit and study reward-related activation within fMRI designs[2,8] and is reliable over time in healthy volunteers[9]. The five cohorts employed a slightly different variation of the MID.

**Discovery cohort**

To examine neural responses to reward anticipation and reward outcome, participants performed a modified version of the MID task[1], in which different cues indicated small and large monetary gains. The task consisted of 66 10-second trials in total and 22 trials per condition. An incentive cue indicating potential rewards was displayed on the left or right side of a black screen for 250 ms on each trial. There were three types of incentive cues (i.e., three within-subject conditions): a circle with two lines (large reward: 10 points), a circle with one line (small reward: 2 points), and a triangle (no reward: 0 points). After a variable delay (4,000-4,500 ms) of fixation on a white crosshair, participants were instructed to respond with a left/right button press as soon as the target appeared. Only responses given within the response interval were considered correct (i.e., reward hit). Using a tracking algorithm, task difficulty (i.e., target duration varied between 100 and 300 ms) was individually adjusted to produce a 66% success rate: the response interval was shortened if the success rate exceeded 66% (making the task more difficult) and lengthened if the success rate was below 66% (making the task easier). Then, feedback on whether and how many points were won during the trial lasted a total of 2000 ms. The inter-trial interval was 3,500–4,150 ms, during which a fixation cross was presented. A trial was considered a hit when the response was made while the target was on the screen. Participants had first completed a practice session outside the scanner (~5 minutes), during which they were instructed that for every 5 points won, they would receive one food snack in the form of small chocolate candies.

**Replication and Clinical cohorts**

The general layout of a trial in this version of the MID task [4] is divided into three phases: anticipation (2000 ms), target (up to 500 ms), and outcome (2000 ms). Immediately prior to scanning, participants were instructed on the task to be performed in the scanner, including being explicitly informed of the meaning of each cue. The target presentation duration was calculated based on individual performance during the training phase to achieve at least a 66% success rate. During anticipation, cue stimuli consisted of three different white geometric shapes: a full circle, representing a chance of gaining 100 points (reward condition), an empty circle, representing a chance of gaining 0 points (control condition), and a full square, representing a chance of losing 100 points (punishment condition). The target was the appearance of a white triangle. Immediately after the response, feedback appeared for 2000 ms, documenting whether the participant had won or lost points as well as their cumulative total at that point. Participants were instructed to respond with a single button press with their right thumb when the target appeared to gain or not lose points. A trial was considered a hit when the response was made while the target was on the screen. The outcome phase provided feedback on money earned from that trial and the total amount earned so far.

**Pharmacological challenge cohorts**

***Single-dose***

This version of the MID task[10] employed four randomized trial types: three active reward-level cues (high (£2), low (£0.20), and control (£0)), and a passive trial that required no response. If the participant pressed the button during the presentation of the fixation cross or within 100 ms of the target presentation (an unrealistic reaction time), the trial would be set as a no-win. Each condition was presented 24 times within a total duration of approximately 13.8 minutes. The three active trial types were conducted within a fixed 10-second window, while the passive trial was a simple 'X' displayed for 4.25 seconds without the fixation, target, or feedback screens displayed in the active trials. Four separate 'playlists' with the 96 trials randomly arranged in each were created, which participants randomly received on each visit according to a Latin square design, to ensure there were no learning effects from completing the same task on each visit. Performance-related criteria were also set to ensure that only data from participants actively and appropriately engaged in the task would have been included in the final analysis. During active trials, a hit was defined as a button press with a reaction time within ±3 standard deviations of the mean, and only participants with a button press rate exceeding 66% in the 500ms response window were included, ensuring active engagement.

***Repeated-dose***

The task contains two trial types (24 win trials and 48 neutral trials)[10]. Participants are instructed to respond as quickly as possible to a target stimulus, and can win money if they respond quickly enough during win trials. The total amount of money to be won is £7.20, with £0 or £0.30 at stake in each trial. Each trial begins with the presentation of a cue stimulus for 500ms, which denotes whether the trial is a win trial (orange square) or a neutral trial (blue square). Following the cue, there is an anticipation period (interstimulus interval (ISI)) which varies randomly between 2, 3, and 4 seconds. The target stimulus (a white square) is then presented for a variable duration (starting at 300ms, 16.67ms subtracted or added each trial depending on performance in previous trial, range 200-400ms), during which time the subject has to respond. The target hit rate was approximately 50%. Following the target, feedback on the outcome of the trial is presented. The duration of feedback is also dynamic, to ensure that the total duration of the target plus the feedback is 1300ms. Following the feedback, an inter-trial interval (ITI) consisting of a fixation point is presented, which varies randomly between 2.2 and 10.2 seconds in one-second increments, on an approximately Poisson distribution4.

**Neuropsychological and clinical assessment**

**Discovery cohort**

The neuropsychological assessment in the Discovery cohort included the Temperament and Character Inventory-Revised (TCI-R)[11,12], which comprised the Novelty Seeking scale and its four temperamental subcomponents (exploratory excitability, impulsiveness, extravagance, and disorderliness) to assess lower order trait dimensions more specifically related to disinhibitory psychopathology, and the Substance Use Risk Profile Scale (SURPS)[13], investigating the role of main personality traits as potential risk factors for addiction and co-morbidity in psychopathology development.

- Specifically, the TCI-R is a self-administered dimensional questionnaire created to evaluate seven basic dimensions of personality as they develop and widely vary within individuals in the general population according to Cloninger's comprehensive biopsychosocial model[12]. Cloninger's personality model includes four temperament dimensions (i.e., Novelty seeking, Harm Avoidance, Reward Dependence, and Persistence) and 3-character dimensions (i.e., Self-directedness, Cooperativeness, and Self-transcendence). In the IMAGEN data collection only the 35 self-reported TCI-R items evaluating the Novelty Seeking, as the tendency to respond impulsively to novel stimuli with active avoidance of frustration, were included to assess lower order trait dimensions more specifically related to disinhibitory psychopathology: participants rated each of the 35 items related to Novelty Seeking on a 5-point Likert scale, with 1='definitely false', 2='mostly false', 3='neither true or false', 4='mostly true', and 5='definitely true'. Thus, we could compute and select for analysis purposes only five personality summary scores, i.e., the four total scores for Exploratory excitability, Impulsiveness, Extravagance and Disorderliness, and the overall Novelty Seeking score to which they contribute as personality sub-dimensions.
- The SURPS is a self-reported questionnaire used mainly in epidemiological and longitudinal designs to investigate the role of 4 main personality traits as potential risk factors for addictive behaviors and co-morbid psychopathology development. Specifically, in the IMAGEN 23-items version of the questionnaire, seven items concur in measuring hopelessness levels (as the tendency to develop bleak expectations about oneself and the future), five items concur in measuring anxiety sensitivity levels (as the fear of anxiety-related physical sensations), five items concur in measuring impulsivity levels (as lack of premeditation and difficulties with response inhibition), and six items concur in measuring sensation seeking levels (as the need for intense and novel experiences). Participants rate each item on a 4-point Likert scale, with 1= 'strongly disagree', 2= 'disagree', 3= 'agree', 4= 'strongly agree'. Thus, we could compute and select four summary scores for analysis purposes, one for each assessed risk personality trait.

To assess cognitive performance across diverse cognitive domains, including executive function, decision-making, and cognitive control, we selected scores from the Passive Avoidance Learning Paradigm (PALP)[14] and from the Cambridge Guessing Task (CGT) included in the Imagen computerized neuropsychological battery (Cambridge Neuropsychological Test Automated Battery – CANTAB; https://www.cambridgecognition.com/cantab/).

- In the PALP, subjects must learn by trial and error to respond to "good" numbers for a monetary reward and withhold response to "bad" numbers to avoid punishment (loss of money). The nine outcome measures in the task were the number of commission errors, denoting too little behavioral inhibition (pressing the button on no-go trials), and the number of omission errors, denoting too much behavioral inhibition (not pressing the button on go trials), as well as the reaction times (ms), across each combination (PP=Punishment-Punishment; RP=Reward-Punishment; RR=Reward-Reward).

- The CGT task was developed to assess decision-making and risk-taking behavior outside a learning context. Relevant information is presented to the subjects "upfront", and there is no need to learn or retrieve information over consecutive trials. On each trial, the subject is presented with a row of ten boxes across the top of the screen, some of which are red and some of which are blue. At the bottom of the screen are rectangles containing the words 'Red' and 'Blue'. The subject must guess whether a yellow token is hidden in a red box or a blue box. In the gambling stages, subjects start with a number of points displayed on the screen and can select a proportion of these points displayed in either rising or falling order in a second box on the screen to gamble on their confidence in this judgment. A stake box on the screen displays the current amount of the bet. The subject must try to accumulate as many points as possible. To make the task shorter and more interesting for adolescents, the time between stakes is reduced from 5s to 2s. Stakes are displayed in ascending order first, and then in descending order. The six scores, including delay aversion, deliberation time, overall proportion bet, quality of decision-making, risk adjustment, and risk-taking, were computed and included in the between-group analysis.

**Replication cohort**

To replicate the neuropsychological and cognitive measures used in the discovery cohort, we employed seven personality scores from the TCI[12] (Novelty Seeking, Harm Avoidance, Reward Dependence, Persistence, Self Directedness, Cooperativeness and Self-Transcendence), four cognitive scores from the Wisconsin Card Sorting test (WCST)[15], which assess executive functions and seven scores from the Continuous Performance Test (CPT)[16], assessing sustained attention and cognitive control. No measure of decision-making and passive avoidance learning was included in the data collection.

- In particular, the WCST is a well-established measure of general executive function evaluating several "frontal" lobe functions, including strategic planning, organized searching, utilizing environmental feedback to shift cognitive sets, directing behavior toward achieving a goal, and modulating impulsive responding. The test uses stimulus and response cards (64-card scoring version) with various forms, colors, and numbers. The participant is told to match the cards, but not how to match them; however, the subject is told whether a particular match is right or wrong. As the test progresses, there are unannounced shifts in the sorting principle, which require the subject to alter her/his approach. Performance data are recorded as the number of completed categories, number of trials, and number of perseverative and non-perseverative errors.
- The CPT–AX examines the attentional domain, specifically sustained and selective attention. During the task, single letters were presented sequentially on the screen, and participants responded when they saw an X after an A. Randomly, a non-cue letter could precede the target (e.g., B-X), or a distractor could follow a cue (e.g., A-Y), or a non-cue letter could precede a distractor (e.g., B-Y). We computed the ratio of the target stimuli actually detected (HIT), the ratio of the false alarms, i.e. the inappropriate responses given to a distractor appearing after the letter "A" (FA)[17] and "B" (FAC), a sensitivity index calculated using the proportion of total false alarms (AP S), the proportion of AY errors

(AY), individual's ability to discriminate signal from noise (DP S), and sensitivity of target detection in relation to the nature of the preceding stimulus (DPC S), to be implemented in the between-group analysis.

**Clinical cohort**

We used the standardized, clinical interview PANSS to assess the severity of positive and negative symptoms before MRI scanning. It comprises 30 items on three subscales: 7 items covering positive symptoms (hallucinations and delusions), 7 covering negative symptoms (blunted affect and social withdrawal), and 16 covering general psychopathology (somatic concern, anxiety, and depression). Each item is scored on an item-specific scale ranging from 1 to 7. The psychometric properties of the PANSS have been well established within adult literature, and it is considered one of the most widely used research measurement tools.

**fMRI Acquisition**

**Discovery cohort**

Structural and functional MRI data were acquired at eight IMAGEN assessment sites with 3-T scanners of different manufacturers (Siemens, Philips, General Electric, and Bruker). The scanning variables were specially chosen to be compatible with all scanners. All sites used the same scanning protocol.

High-resolution T1-weighted 3D structural images were acquired for anatomical localization and coregistration with the functional time series. Blood-oxygen-level-dependent (BOLD) functional images were acquired with a gradient-echo echoplanar imaging (EPI) sequence and using a relatively short echo time to optimize reliable imaging of subcortical areas. For each participant, 300 volumes were acquired, and each volume consisted of 40 slices aligned to the anterior commissure/posterior commissure line (2.4 mm slice thickness and 1 mm gap). The echo time was optimized (echo time=30 ms; repetition time=2200 ms) to provide reliable imaging of subcortical areas. The acquisition parameters per site and quality control procedures are described elsewhere[1].

**Replication cohort and Clinical cohort**

Structural and functional MRI data were acquired with a 3 T Philips Ingenia scanner. High-resolution T1-weighted 3D structural images were acquired for anatomical localization and coregistration with the functional time series. BOLD functional images were acquired with gradient-recall echo-planar imaging with the following parameters: TR = 2000 ms; TE = 38 ms; flip angle = 90; 64 × 64 matrices; FOV = 240 mm; and 38 3.6 mm slices acquired with an interleaved order of slice acquisition.

**Pharmacological challenge cohorts**

***Single-dose***

All scans were conducted on a GE MR750 3-Tesla scanner using a 12-channel receive-only head coil. Functional scans (MID and breath-hold) were carried out using a temporal series of Gradient Recalled Echo Planar Imaging (GE-EPI) whole brain scans, each comprising 38 near-axial slices, with an isotropic spatial resolution of 3.3 mm and the following parameters: TR=2000 ms; TE=28 ms; flip angle=75; number of volumes=414; FOV=214 mm.

***Repeated-dose***

Structural and functional MRI images were acquired on a 3T Siemens Magnetom Prisma with a 64- channel head coil. High resolution T1 weighted volumes were acquired using a MP2RAGE sequence (176 slices, FOV read = 256 mm, TR = 5000.0 ms, TE = 2.98 ms, TI 1 = 707 ms, TI 2 = 2500 ms, flip angle 1 = 4°, flip angle 2 = 5°, 1 mm isotropic voxels, bandwidth = 240 Hz/pixel, echo spacing = 7.1 ms). For the MID task, at each scanning session, 304 functional volumes were acquired using a BOLD echo planar imaging sequence (FOV read 220mm, 42 slices, slice thickness 3 mm, voxel size 3.4 x 3.4 x 3mm, TR 2400 ms, TE 30 ms). The first six volumes of each functional run were discarded to allow for T1 saturation effects.

**fMRI Processing and task-activity modeling**

We followed the same preprocessing pipeline for the Discovery, the independent Replication, and the Clinical cohorts using the open-source containerized Harmonized AnaLysis of Functional MRI pipeline (HALFpipe) version 1.2.2[18] built using fMRIprep version 20.2.7[19] in combination with the Magnetic Resonance Imaging Quality Control tool (MRIQC)[3]. Consensus steps for structural images included skull stripping, tissue segmentation, and spatial normalization. For functional images, pre-processing included motion correction (and motion parameter extraction), coregistration, spatial normalization to MNI152 NLIN 2009c (asymmetric) space, and reslicing to 2mm3 and smoothing with a 6 mm FWHM Gaussian kernel. Data were high-pass filtered with a cutoff of 125 s to remove low-frequency drifts. Grand mean scaling was applied to the data. Participants were excluded if the mean root-mean-square framewise displacement exceeded 0.5mm. MRIQC provides an interactive widget to rate the quality of each image[19] for each site.

A first-level general linear model (GLM) was run to model BOLD responses to events of interest separately for each subject. GLM regressors describing the stimulus presentations for each experimental condition (i.e., cues by type, successful/unsuccessful outcomes, missing and error trials) are convolved with a double Gamma HRF, and the overall model is fit for each voxel in the brain using FSL.

Our contrasts of interest were created to capture the individual reward sensitivity by contrasting the BOLD signal of rewarded trials to control cue events. Thus, for the anticipation phase, we contrasted the brain activation corresponding to the successful trials preceded by reward cues to the brain activation corresponding to the successful trials preceded by a control cue (Discovery cohort: Hit Large Reward > No-Reward; independent Replication cohort: Reward > No-Reward). Similarly, for the outcome phase, we contrasted the brain activation during the appearance of the successful feedback preceded by a reward cue during anticipation to the brain activation during the appearance of the successful feedback preceded by a control cue (Discovery cohort: Hit Large Reward > No-Reward; independent Replication cohort: Reward > No-Reward).

**Pharmacological challenge cohorts**

***Single-dose***

Preprocessing was conducted in the Statistical Parametric Mapping (SPM) analysis suite, version 12, on Matlab 8.2.0.701, and included resetting of image origins, slice time correction, two-pass realignment, co-registration, and normalization to MNI space using DARTEL (Diffeomorphic anatomical registration through exponentiated lie algebra)[20], and smoothing using an 8 mm FWHM kernel. The origin of the functional and structural images was reset to the anterior commissure-posterior commissure line. Functional images were slice time corrected (reference slice: 19). An initial between-session alignment was performed for each participant, where the first volume of sessions two and three was aligned to the first volume of the first session, prior to two-pass realignment within each session. All volumes were then realigned to the mean image of all three sessions. The T1-weighted image for each subject was then coregistered to the resampled mean functional image from the realignment step, using the normalized mutual information objective function in SPM, and a DARTEL[20] template created from the T1-w images. The realigned and coregistered functional volumes were resliced to original voxel sizes, and the DARTEL flow fields were applied to warp the data into MNI space. Normalized images were smoothed using an 8mm FWHM kernel. Motion and framewise displacement parameters estimated during the realignment process were added as regressors in the first-level design matrix. Any volumes with a displacement of 1mm or more were flagged and marked with a 3-TR regressor (to include the volumes on either side) in the first-level design matrix. Any scan that required more than 10% of the volumes of the full run being regressed out in this manner resulted in that participant being removed from the analysis. The realignment parameters were visually inspected, and any time series for which the maximum detected translation from the first volume was greater than the dimensions of one voxel, or those that indicated stimulus-correlated movement, were flagged for exclusion. Three participants were excluded based on these criteria. These participants were identified and excluded in parallel to data collection, allowing them to be replaced by new recruits to maintain a suitably powered study. Participants excluded due to head motion do not form part of the sample described above.

Contrasts of interest were set to explore the main effect of anticipation of reward (High Cue > Neutral Cue) and the main effect of receipt of reward (High Win > Neutral Cue). The MID was modelled as outlined elsewhere[21]. Three cue regressors (high win, low win, and neutral) were defined for the anticipatory period depending on the cue presented (as a result of temporal jittering; this period had a variable total amount of time between 4050ms and 4400ms). The target was defined by a single regressor of 500ms. Five feedback regressors (high win, low win, high no win, low no win, neutral feedback) were defined for the feedback period, depending on the cue type and outcome (win or no win), and set for a fixed period (1450ms). The entire duration of passive trials was defined as a single event of 4250ms. Motion and framewise displacement parameters estimated during the realignment process were also added, resulting in the model consisting of the ten task regressors above and seven movement-related parameters. Performance-related criteria were also set to ensure that only data from participants who were actively and appropriately engaged in the task were included in the final analysis.

***Repeated-dose***

Image processing was performed using FSL version 6.00. Anatomical images were preprocessed using the FSL_anat function in FSL to perform bias correction, transformation into a standard stereotactic space (MNI152) and brain extraction using the BET extraction tool. Functional image series were pre-processed with a 100s high-pass filter, head motion correction, 6mm full-width at half maximum spatial smoothing, and co-registration to the T1-weighted structural image before transformation to standard space. The times of the stimulus conditions were convolved with a gamma function to simulate the haemodynamic response function. Head-motion parameters were included in the first-level (subject-level) models as nuisance regressors. Temporal derivatives of task-related events and neutral hit and neutral miss explanatory variables were also included as nuisance regressors. The contrasts of interest in the MID task were the subtraction contrasts of the reward anticipation period minus the neutral anticipation period (reward anticipation), and the reward outcome period minus the missed reward outcome period (reward outcome). We conducted a second level analysis to calculate the group mean of all baseline scans (regardless of subsequent treatment order or condition) to ensure that the tasks were activating the reward network. We used FSL's FLAME 1 model, thresholded at $Z > 3.1$, using a corrected cluster significance threshold of $p = 0.05$.

**fMRI trial-by-trial analysis**

**Replication cohort**

To investigate group differences in reward anticipation between sign-trackers (ST) and goal-trackers (GT), we analyzed trial-by-trial variability across 28 reward and 28 no-reward trials in the independent Replication cohort. We employed AFNI's *3dDeconvolve* tool[22] to perform deconvolution analyses, obtaining single-trial beta estimates via the beta-series regression method[23].

Specifically, the *-stim_times_IM* option was used within *3dDeconvolve* to model each trial individually for both reward and no-reward conditions. This approach assigns a unique regressor to each trial, enabling the extraction of distinct beta coefficients that reflect the hemodynamic response associated with each event. The resulting beta estimates were then used to assess trial-by-trial variability and compare neural responses between ST and GT during reward anticipation.

**Statistical Analyses**

Based on a recent meta-analysis[24] on the MID task—which identified a set of regions exhibiting enhanced BOLD activity during both reward anticipation (Reward vs. No Reward) and reward outcome (Reward vs. No Reward)—a total of 22 regions of interest (ROIs) was defined for our study. None of the 81 studies involved the IMAGEN data employed in this study, i.e., follow-up 3. Specifically, for all four cohorts, we created 12 ROIs for the anticipation phase (including bilateral ventral striatum, thalamus, supplementary motor area, bilateral motor cortex, bilateral anterior insula, bilateral lateral occipital cortex, and bilateral middle frontal gyrus) and 10 ROIs for the outcome phase (including bilateral ventral striatum, ventromedial prefrontal cortex, bilateral lateral occipital cortex, posterior cingulate cortex, bilateral superior parietal lobule, left middle frontal gyrus, and right inferior frontal gyrus), showed in Supplementary Fig. 1. We then extracted the individual signal from the anticipation and outcome ROIs in each individual activation map, grouped by the task phase, using an uncorrected threshold of $p<.001$ through MarsBar implemented in SPM12 (http://www.fil.ion.ucl.ac.uk/spm).

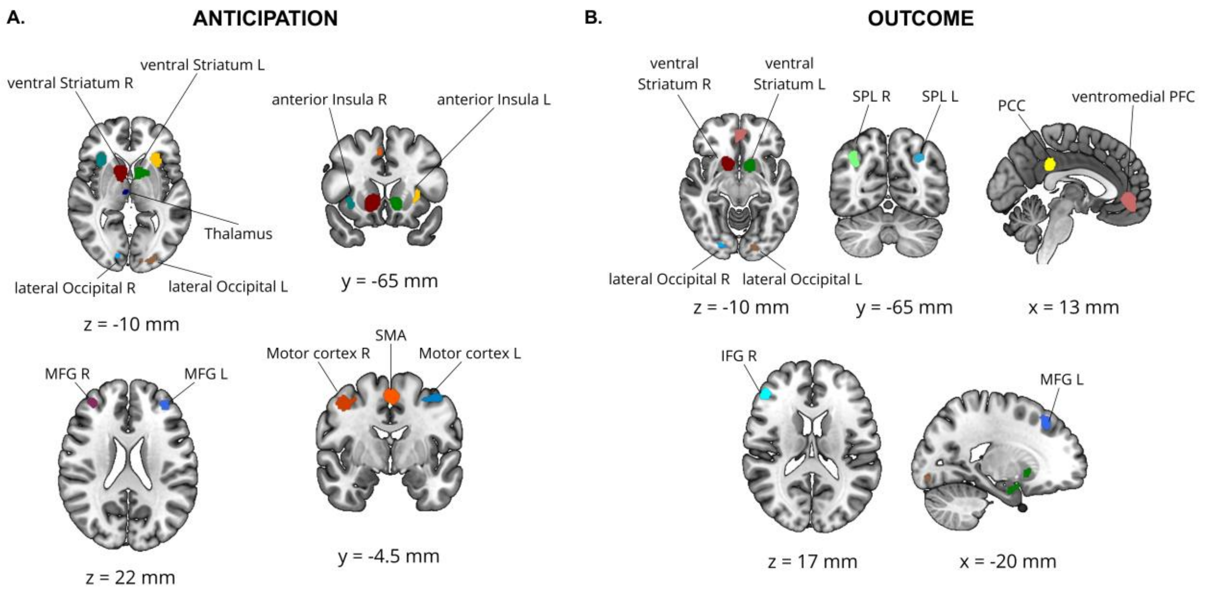


**Supplementary Fig. 1. Regions of interest (ROIs) employed in the current study, derived from meta-analytic maps.** (**A**) Twelve ROIs were utilized for feature extraction of individual BOLD signals during the anticipation phase of the Monetary Incentive Delay (MID) task. (**B**) Ten ROIs were used for feature extraction during the outcome phase of the MID task. *Abbreviations:* ROIs=regions of interest; BOLD=blood-oxygen-level-dependent; MID=monetary incentive delay task; R=right; L=left; MFG=middle frontal gyrus; SMA=supplementary motor area; SPL=superior parietal lobe; PCC=posterior cingulate cortex; PFC=prefrontal cortex; IFG=inferior frontal gyrus.

#### Cluster Analysis

##### Discovery cohort

To test our hypothesis that individuals can be grouped by their brain activation during the performance of a reward task, we performed a hierarchical K-means clustering[25] using the *hkmeans()* R function in *factoextra* package by including as features the BOLD signal intensity values extracted from the ROIs on the individual task-related activity during i) the anticipation and ii) the outcome phases during the MID task. The features were initially checked for the potential presence of outliers using Rosner's generalized extreme Studentized deviate test[26] from the *EnvStats* R package ($p<0.05$). It is indicated for up to *k* potential outliers in a large dataset, assuming the data without any outliers comes from a normal (Gaussian) distribution. It removes one potential outlier at a time, recalculates test statistics, and continues until the specified number of potential outliers is evaluated, making it useful for datasets where multiple anomalies may be present. Then, values were scaled to a -1 – 1 range to remove the differences between features, and we adjusted the data for the effect of the site with the function *adjust()* from the *datawizard* R package, which returned residuals of the regression models.

We determined the optimal number of clusters ($k$ = 3) by combining two complementary approaches: the *NbClust* R package and a hypothesis-driven approach. The latter is based on prior work on the ST/GT model[12,81], yielding the ST, GT, and INT phenotypes. Formally, NbClust evaluated 30 established cluster validity indices across a leave-site-out validation framework and favored a k = 2 solution, receiving the highest number of votes across indices (Supplementary Fig. 2, panel A). Despite this, we adopted k = 3 for the Discovery cohort because it aligns with the well-established three-group distribution in the animal literature and because, across individual sites, k = 3 was frequently favored. This choice was deliberate and hypothesis-driven: by extracting only the ST and GT centroids for projection onto independent cohorts, the INT group serves as a buffer zone, reducing noise and maximizing phenotypic separation. As visible in Supplementary Fig. 2, panel B, when the k = 3 solution is visualized in PCA space, the INT group occupies the region between the ST and GT confidence ellipses, with its centroid at the midpoint along Dim1, while the ST and GT ellipses remain well separated.

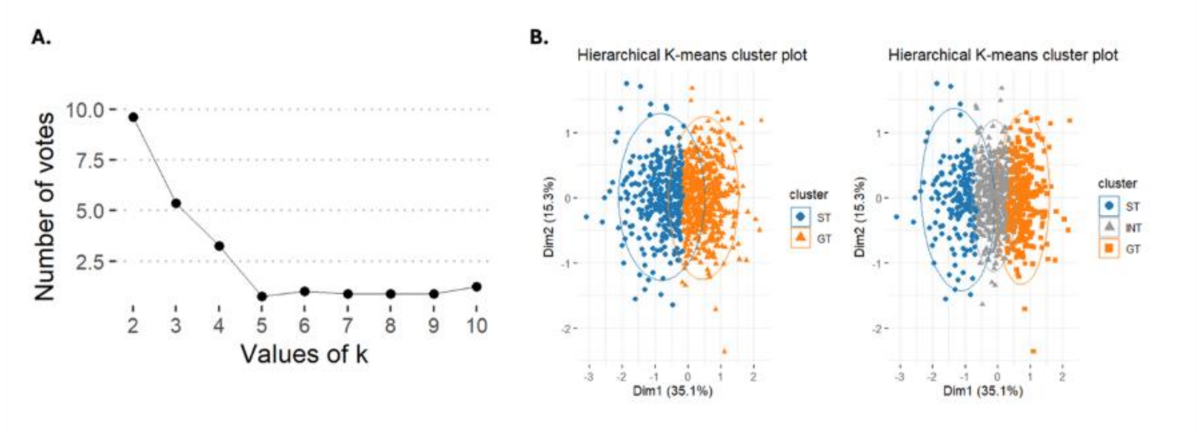


**Supplementary Fig. 2. (A) Elbow plot.** The elbow plot in the left panel shows the total within-cluster sum of squares (WSS) as a function of the number of clusters (k), with a pronounced inflection at k = 2, indicating that two clusters provide the optimal partition of the data. **(B) Hierarchical k-means clustering solution visualized on the first two principal components.** On the left, two-cluster solution (k = 2) derived from hierarchical k-means clustering, identifying Sign Trackers (ST) and Goal-Trackers (GT). On the right, three-cluster solution (k = 3), revealing an intermediate group (INT) positioned between the ST and GT clusters. Each point represents an individual participant projected onto the first two principal components (Dim1 and Dim2), explaining 35.1% and 15.3% of the variance, respectively. Ellipses indicate the dispersion of observations within each cluster. Colors denote cluster membership.

This is also supported by the GMM-based BIC analysis, where BIC favored two components and increased monotonically beyond k = 2, providing no evidence for finer subgroupings (Supplementary Fig. 3, panel A). Specifically, we conducted three independent analyses to test whether the data are better described as a continuum or as separable categories. If a single latent dimension were sufficient, GMM-based model selection using BIC would have favored k=1. Instead, BIC formally favored a two-component solution (ΔBIC = 17.9), indicating that the multivariate neural data across 22 ROIs are better described by two partially overlapping distributions than by a single Gaussian (Supplementary Fig. 3, panel A). Ashman's D on the first principal component of this space was 2.62, exceeding the conventional bimodality threshold of 2.0[27] (Supplementary Fig. 3, panel B). We note that PC1 explains 35.1% of the total variance, and the partial overlap visible in its density plot reflects the compression of a 22-dimensional structure onto a single axis rather than the true degree of separation between groups. Consistent with this, the Mahalanobis distance between ST and GT centroids computed across all 22 dimensions was 2.77, exceeding the permutation-derived null by a factor of 8.5 ($p < 0.001$; Supplementary Fig. 3, panel C). Within this two-component structure, the in-between participants are best regarded as the less prototypical members of the two distributions, occupying the region where the ST and GT distributions overlap rather than forming a discrete third group [28,29]. The convergence of these analyses supports treating ST and GT as statistically separable profiles rather than arbitrary partitions of a continuum.

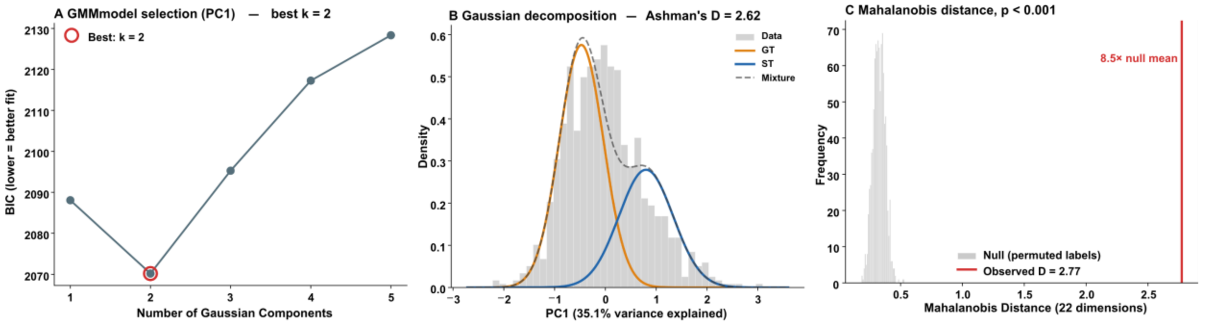


**Supplementary Fig. 3. Statistical Evidence for Categorical Separation of ST and GT Phenotypes. (A) GMM Model Selection:** Bayesian Information Criterion (BIC) values for Gaussian Mixture Models with 1 to 5 components. The minimum BIC at k=2 (Delta BIC = 17.9 vs. k=1) indicates that two partially overlapping distributions provide a superior fit compared to a single Gaussian continuum. **(B) Gaussian Decomposition:** Density plot of the first principal component (PC1, 35.1% variance) showing the distribution of the two identified components. Ashman's D = 2.62 exceeds the conventional threshold (D > 2.0) for significant bimodality. (**C) Multivariate Distance:** Mahalanobis distance between ST and GT centroids computed across all 22 ROI dimensions. The observed distance (D = 2.77) is 8.5 times larger than the mean of the permutation-derived null distribution (grey histogram), confirming highly significant spatial separation in the multivariate neural space ($p < 0.001$).

In our clustering, we hypothesized to find similar groups based on brain activity during the two separate phases of the task. Specifically, ST with elevated brain activity during the anticipation phase, GT with elevated brain activity during the outcome phase, and intermediates with mixed brain activity during both phases.

To evaluate the potential effect of sex in clustering, we performed a hierarchical k-means clustering (k=3), separated for males (N=434) and females (N=454), replicating the results obtained with the clustering applied on the full dataset. Results are shown in Supplementary Fig. 4.

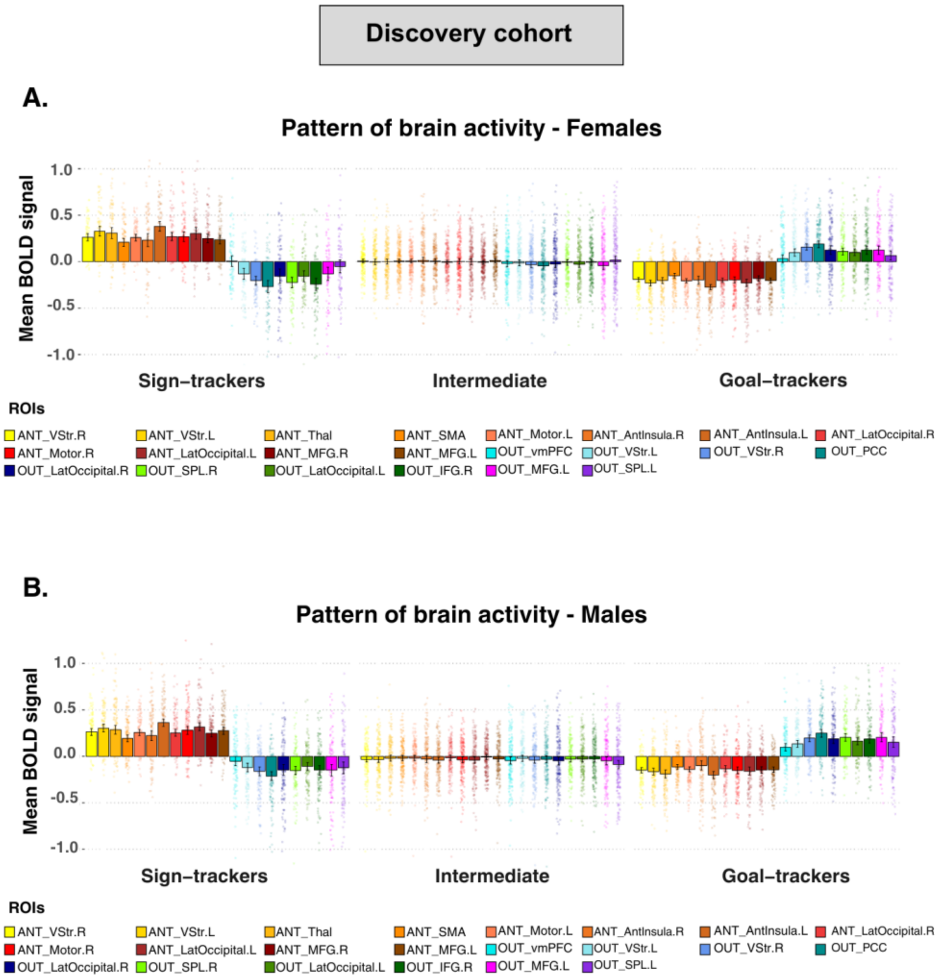


**Supplementary Fig. 4. Pattern of brain activity grouped by sex in the Discovery cohort.** Barplots showing BOLD activity in each ROI for each cluster, i.e., sign-trackers, intermediate, and goal-trackers, in females (**A**) and males (**B**) in the Discovery cohort. *Abbreviations:* ANT=anticipation; OUT=outcome; R=right; L=left; VStr=ventral striatum; SPL=superior parietal lobule; Thal=thalamus; MFG=middle frontal gyrus; SMA=supplementary motor area; IFG=inferior frontal gyrus; vmPFC=ventromedial prefrontal cortex; PCC=posterior cingulate cortex.

#### Leave-Site-Out Validation

To validate the clustering performed on the full dataset (N=890) described above, we performed the hierarchical k-means clustering[25] within a leave-site-out validation framework iteratively for each site. To this end, the dataset was initially divided into a training set (seven sites) and a testing set (one site). Scaling (-1 to 1 range) was applied separately in both sets, and residuals of regression models for the effect of the site were implemented only in the training set. The hierarchical k-means clustering[25] was performed on the training set with $k$=3 using Euclidean distance and Ward's linkage. To validate this algorithm on the testing set, we assigned the cluster membership by computing the Euclidean distance between the testing set and the two training cluster centers, corresponding to the sign and goal-trackers, due to the small sample size of the testing set.

We then computed the clustering on the testing set to label the testing set data. We calculated a confusion matrix from the *caret* R package to evaluate the cross-tabulation of the cluster assignments with associated statistics, i.e., sensitivity, specificity, balanced, and overall accuracy. For two-class problems, the sensitivity, specificity, positive predictive value, and negative predictive value are calculated using the *positive* argument (if there are only two factor levels, the first level will be used as the "positive" result). The overall accuracy rate is computed along with a 95 percent confidence interval for this rate and a one-sided test to see if the accuracy is better than the "no information rate", which is taken to be the largest class percentage in the data. Balanced accuracy is computed in relation to sensitivity and specificity and, in this study, is used as a measure of replicability. To overcome the issue related to the incorrect order, we previously realigned the levels with *solve_LSAP()* function from the *clue* R package[30].

**Replication cohort**

To assess the generalizability of our neurophysiological-based clustering, we applied the Discovery clustering algorithm to our independent Replication cohort. As in the Discovery cohort, the BOLD signal was scaled (-1 to 1 range) to remove differences between features across samples. To assess the goodness of the Discovery algorithm in identifying the same groups in an independent dataset, everyone in the Replication dataset was assigned to a cluster based on its proximity to the Discovery cluster centers computed on the full dataset. Due to the small sample size, we projected only the two centroids – ST and GT – to obtain predicted clusters and then we performed a hierarchical k-means clustering[25] on the Replication dataset (*k*=2) with Euclidean distance and Ward's linkage to obtain the actual data-driven cluster memberships. We compared the predicted clusters with the actual clusters through a confusion matrix, obtaining overall accuracy and the statistical measures of the sensitivity (true positive rates), specificity (true negative rates), and replicability for the two classes.

**Association with personality and cognitive measures**

**Discovery cohort**

Each variable was scaled (0-1 range) to remove the differences between features and to remove any confounding effects due to the site. We used the residuals of the regression models computed with the function *adjust()*. After removing outliers with the Rosner test[26], we separately evaluated the differences between the ST and GT across all the neuropsychological measures through the Wilcoxon rank-sum test. To determine effect size (r equivalent), we used the *rstatix* R package[31] by dividing the Wilcoxon Z-transformed values by the square root of the number of observations[32]. All significant p-values were <0.05, false discovery rate (FDR)-corrected [33]. Additionally, we conducted the same analysis with ST and GT obtained from a hierarchical k-means clustering with k=2 to avoid the loss of subjects in the intermediate group that may undermine the statistical power.

**Replication cohort**

As in the Discovery cohort, we applied a 0-1 range scaling to each variable to remove the differences between features. After removing outliers with the Rosner test[26], we performed the Wilcoxon rank-sum test ($p_{FDR}$<0.05, one-tailed) to evaluate the differences between clusters across all the neuropsychological measures. We excluded subjects that were misidentified in the cross-tabulation of cluster assignments from these analyses to reduce noise in the data (N=202). To replicate PALP results, showing ST more efficient than GT in learning shifted contingencies, we used a one-tailed Wilcoxon rank-sum test to evaluate the specific hypothesis that ST would perform better than GT at the WCST. To determine effect size (r equivalent), we used the *rstatix* R package[31] by dividing the Wilcoxon Z-transformed values by the square root of the number of observations[32]. Additionally, we conducted the same analysis with ST and GT obtained from a hierarchical k-means clustering with k=2 to avoid the loss of the mis-identified subjects that may undermine the statistical power.

**Trial-by-trial Analysis**

Trial-by-trial analyses were conducted using the beta-series method[23] to assess whether ST and GT differentially attributed incentive salience to reward anticipation cues[34]. For this reason, we used a stronger ventral striatum response over trials as an index of incentive salience. As we predicted these trial-by-trial group differences would be specific to the reward condition, we modeled reward and no-reward trials in separate analyses. To predict trial-by-trial ventral-striatal BOLD responses, we used a linear mixed-effects model (LME) with fixed effects for Group, Trial, and their interaction. The models included only individuals correctly identified as GT or ST. Before entering the linear mixed model analysis, the dependent variables were winsorized to reduce the influence of outliers while retaining all data points. Specifically, values below the first percentile and above the 99th percentile were winsorized, a process known as 2% winsorization. Sex, age, and quadratic age effects were included as covariates to account for potential non-linear age-related differences in brain activity. Following best practices[35], we initially attempted to fit a maximal random effects structure with by-subject random intercepts and by-subject random slopes for Trial, including the correlation between them. As this model failed to converge, we implemented a more parsimonious no-random-correlations (NRC) model that excluded the correlation parameter. This is a recommended approach to achieve a stable solution in such cases while retaining the critical random effect variance. All models were fitted using REML (Restricted Maximum Likelihood).

**Group-level task activity across clusters**

In both the Discovery and Replication samples, as well as in the Clinical cohort, to evaluate task-related activity at the group level for each cluster, i.e., ST and GT, we performed separate one-sample t-tests for each task phase by entering the individual contrast maps described above using SPM12. All statistics were non-parametrically corrected for multiple comparisons through the Threshold-Free Cluster Enhancement approach (TFCE)[36]. Significance was set at $\alpha$ <0.05, using the family-wise error-rate (FWE) for multiple comparison correction.

**Pharmacological challenge cohorts**

***Single-dose***

**BOLD activity in the anticipation and outcome ROIs**

To assess the effect of the drug on BOLD activity, preliminary analyses were conducted to determine whether the BOLD signal extracted from anticipation and outcome ROIs could be combined to generate a single estimate for each reward phase. Two approaches were employed for this purpose. The first approach involved computing the mean BOLD signal separately for anticipation and outcome, yielding two distinct estimates corresponding to the respective reward phases. The second approach standardized and linearly combined the BOLD signal from anticipation and outcome ROIs into a single principal component (PC) for each phase. Given the high correlation between the mean-based and PC-based estimates (Anticipation: r=.999; Outcome: r=.987), subsequent analyses in this cohort were conducted using the mean-based estimates.

**Placebo vs. Drug**

A linear mixed-effects model (LMM) was implemented to investigate the effects of a D2 antagonist dose on reward-related responses during both the anticipation and outcome phases of the MID. The analysis was performed using R (version 2023.09.1+494), employing the *lme4* package for model fitting and *lmerTest* for significance testing. The *clubSandwich* package was used to compute robust standard errors to account for potential heteroskedasticity and violations of model assumptions, ensuring valid standard errors under model misspecification. Robust standard errors were computed using a heteroskedasticity-consistent (HC) covariance matrix estimator, specifically CR2, which adjusts for within-subject correlations and potential non-sphericity in the data.

The dependent variable used in the model was the BOLD signal, averaged over the anticipation ROIs or the outcome ROIs. The primary model included reward phase (anticipation vs. outcome), drug (placebo vs. drug), and cluster (Sign-Trackers [ST] vs. Goal-Trackers [GT]) as fixed effects, along with their full factorial interactions. Age was included as a second-degree polynomial term to model potential nonlinear effects. Individual differences in risperidone or haloperidol levels were accounted for using a binomial variable as a covariate. To account for repeated measures, a random intercept was included for each participant.

Model assumptions were checked using the *performance* package in *R*, examining normality of residuals, homoscedasticity, and potential influential data points. Likelihood ratio tests were performed using drop1 to assess whether the inclusion of higher-order interactions significantly improved model fit. Given that a significant three-way interaction was observed, two additional models were fitted separately for anticipation and outcome phases, using the same specifications described above.

To follow up on significant interactions, estimated marginal means (EMMs) were computed using the *emmeans* package in *R*. Post-hoc comparisons were performed for the effect of drug within each cluster, the difference between ST and GT within each drug condition, and the interaction between drug and phase within each cluster. To control for multiple comparisons, Tukey's method was applied, providing strong control over family-wise error rate while maintaining adequate power for pairwise contrasts.

**Dose-response effect of risperidone during the reward anticipation phase**

Since the Placebo vs. Drug analysis showed an interaction between cluster and drug only for the reward anticipation condition, the dose-response effect was tested only on this reward phase in the 17 subjects included in the risperidone study. An LMM was used to investigate the effects of placebo, low-dose risperidone, and high-dose risperidone on response values during the anticipation phase of a reward-related task. In the LMM analysis, drug condition was treated as a categorical fixed effect with three levels (placebo, low-dose risperidone, high-dose risperidone), cluster (Sign-Trackers [ST] vs. Goal-Trackers [GT]) was treated as a fixed effect factor, age was modeled as a second-degree polynomial, and participants were included as a random effect to account for repeated measures within subjects.

Given the three-level nature of the drug factor, different model specifications were tested to determine both categorical and continuous dose-response relationships, including linear and quadratic trends. In addition to the categorical model used in the primary analysis, to further explore potential dose-dependent effects, additional models were fitted, including an ordinal model testing for a linear trend across drug conditions, a quadratic model testing for a curvilinear relationship, and a stepwise model testing the contrast between high-dose risperidone and all other conditions combined. Model selection was performed using likelihood ratio tests (LRTs) comparing each model against the null model, with significance assessed using Chi-square tests. Post-hoc analyses were conducted using EMMs, with pairwise comparisons between drug conditions adjusted using Tukey's method and FDR corrections.

**Subjective drug effects**

The psychometric properties of the visual analogue scale (VAS) items were evaluated in a two-step process using *RStudio*. First, an Exploratory Factor Analysis (EFA) was conducted on the 16 original items from the Placebo Pre-dose condition to identify the underlying factor structure in a baseline state, free from any pharmacological intervention. Following standard practices, we used maximum likelihood estimation with an oblimin (oblique) rotation to allow for correlated factors. The number of factors to retain was determined by a convergence of criteria: the Kaiser criterion (eigenvalues > 1), a scree plot inflection point, and parallel analysis based on 100 permutations.

Second, based on the EFA results and an iterative process of model refinement to improve fit and parsimony, a 10-item, 3-factor model was specified for Confirmatory Factor Analysis (CFA). This refined model was composed of Factor 1 (Energy), which included the items Alert_to_Drowsy (loading=0.85), Strong_to_Feeble (loading=0.89), and WellCoordinated_to_Clumsy (loading=0.92); Factor 2 (Calm), comprising Antagonistic_to_Friendly (loading=0.97), Troubled_to_Tranquil (loading=0.82), and Tense_to_Relaxed (loading=0.63); and Factor 3 (Focus), consisting of MentallySlow_to_QuickWitted (loading=0.90), Incompetent_to_Proficient (loading=0.92), Attentive_to_Dreamy (loading=-0.59), and Muzzy_to_ClearHeaded (loading=0.92). The observed loadings are high, indicating strong associations between items and their respective factors. The score of Factor 1 was reversed to maintain a consistent scaling of low to high score. Notably, our 3-factor solution exhibits both similarities and distinctions compared to foundational factor analyses of similar visual analogue scales from the 1970s. Bond and Lader (1974)[37] identified three factors: 'alertness', 'contentedness', and 'calmness'. Their 'alertness' factor encompassed items related to mental clarity and physical vigor, aligning with items in both our Energy and Focus factors. Their 'calmness' factor (specifically comprising 'Calm' and 'Relaxed' items) strongly corresponds to our Calm factor. Herbert et al. (1976)[38] identified two primary factors, 'alertness' and 'tranquillity'. His 'alertness' factor similarly combined aspects of mental and psychomotor performance, while the 'tranquillity' factor merged elements of contentedness and calmness. Our model's differentiation of the broader historical 'alertness' domain into distinct Energy and Focus factors provides a more granular perspective on these dimensions.

The adequacy of this 3-factor structure was formally tested using the *lavaan* package. The model was fit separately to the data from four distinct conditions: placebo predose, 90 minutes post-dose in the placebo condition, predose in the drug condition, and 90 minutes post-dose in the drug condition. Model fitting was assessed using standard indices, including the Comparative Fit Index (CFI), Tucker-Lewis Index (TLI), Root Mean Square Error of Approximation (RMSEA), and Standardized Root Mean Square Residual (SRMR). To justify the comparison of factor scores across conditions, a multi-group CFA was performed to test for measurement invariance (configural, metric, scalar, and strict) across the three conditions that were not used in the EFA. Finally, for each participant at each timepoint, factor scores for Energy, Calm, and Focus were estimated and saved using the *lavPredict* function for subsequent analysis. As an ultimate step, factor scores for Energy were reversed so that higher values consistently reflected greater energy.

The derived factor scores were analyzed using LMMs to investigate the effects of Time (prepose, post-dose), Cluster (GT, ST), and Phase (Placebo, Drug). Analyses were conducted in R using the *lme4* and *lmerTest* packages. To align with the study design, separate analyses were conducted for the Placebo and Drug phases. For each of the three factors within each phase, an LMM was fitted with the factor score as the dependent variable. In the applied model, cluster, Time, and their interaction were included as fixed effects to test our primary hypothesis. Participant age, its quadratic term, and drug cohort (risperidone, haloperidol) were included as covariates. A random intercept for each subject was included to account for the non-independence of repeated measures. P-values for the fixed effects were obtained using Satterthwaite's degrees of freedom approximation. To control for multiple comparisons across the three factors within each phase, the p-values for the Cluster * Time interaction terms were adjusted using the FDR correction. Significant interactions were followed up with post-hoc pairwise comparisons using a Tukey correction for multiple comparisons.

The final 10-item, 3-factor model demonstrated an improved and adequate fit for the data compared to the initial 16-item EFA solution. In the Predose condition, fit indices were acceptable. Fit remained adequate across the other conditions, with CFI values of .886 in the predose condition during drug and .858 for the post-dose condition during drug, and SRMR < .09 in all conditions. Because of the small sample size, RMSEA values were elevated (> .21). Multi-group CFA supported measurement invariance across the post-dose condition during placebo, the pre-dose condition during drug, and the post-dose condition during drug. The criteria for configural, metric ($\Delta$CFI=.003, p=.223), and scalar invariance ($\Delta$CFI=.002, p=.253) were met, indicating that the factor structure, item loadings, and item intercepts were equivalent across conditions. This justifies the direct comparison of latent factor scores. Strict invariance was not supported, suggesting that residual variances differed between conditions.

### RESULTS

**Leave-Site-Out Validation**

**Discovery cohort**

Despite inter-site variability (Supplementary Fig. 5), the leave-site-out (LSO) cross-validation analysis to assess cluster stability produced an overall accuracy of 86.7% and a measure of replicability of 84.9% (SD=9%), based on the average between sensitivity (87.8%) and specificity (82%).

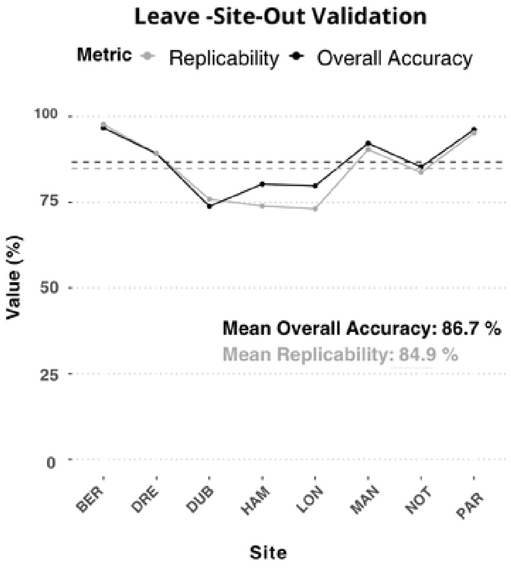


**Supplementary Fig. 5. Leave-site-out validation. In the Discovery multi-site cohort, t**he clusters were highly reproducible across eight imaging sites (86.7% accuracy, 84.9% replicability).

**Association with personality and cognitive measures**

Despite larger sample size, when repeating these analyses using a k = 2 solution, which directly identifies ST and GT without an intermediate group, in the Discovery cohort identical neuropsychological results were obtained in the PALP. On the contrary, we did not find any significant difference regarding the SWM and the CGT (Supplementary Fig. 6, left panel). As in Discovery, in our Replication cohort, when using a k = 2 solution, which directly identifies ST and GT, we did not find any significant difference in the WCST (Supplementary Fig. 6, right panel).

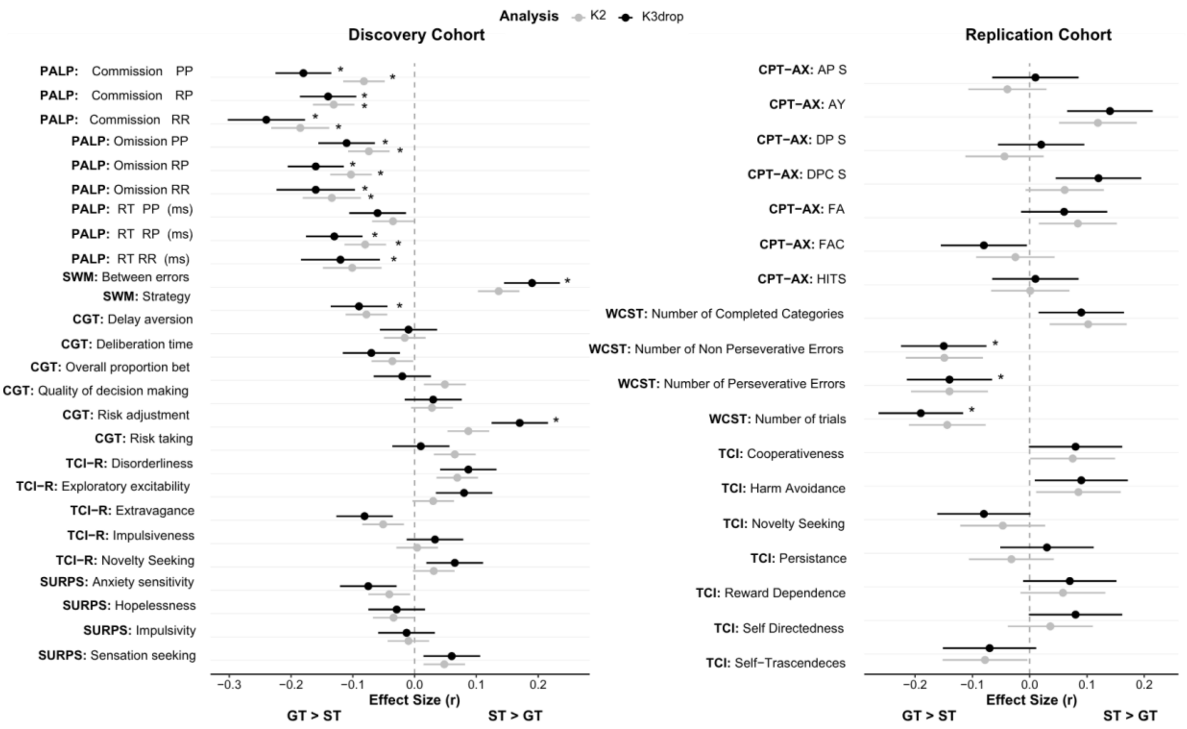


**Supplementary Fig. 6. Stability of Cognitive and Personality Profiling across Clustering Methods.** Comparison of effect sizes (*r*) derived from Wilcoxon rank-sum tests for associations between cluster membership (GT vs. ST) and neuropsychological measures, using two different analytical approaches: K2 (grey) and K3drop (black). **Left:** Discovery cohort results covering Reward Sensitivity (PALP), Cognition (SWM, CGT), and Personality (TCI-R, SURPS). **Right:** Replication cohort results for Cognition (CPT-AX, WCST) and Personality (TCI). Positive values indicate higher scores in ST, while negative values indicate higher scores in GT. The consistency between K2 and K3drop results underscores the robustness of the behavioral phenotypes ($p_{FDR} < 0.05$).

##### Robustness to the width of the intermediate group

To verify that the neuropsychological dissociation in the Discovery cohort did not arise from the exclusion of the intermediate group, we repeated the comparisons with ST and GT defined by a percentile criterion on the first principal component of the reward-related features rather than by clustering. At three thresholds we classified the most ST-like and most GT-like 45%, 40%, and 30% of participants as the two phenotypes and treated the remaining 10%, 20%, and 40% as intermediate, yielding analytic samples of N=392 ST and N=396 GT (10% intermediate), N=349 and N=351 (20%), and N=260 and N=262 (40%). Two-sided Wilcoxon rank-sum tests were computed at each threshold, with effect sizes (r) obtained as the Z-transformed statistic divided by the square root of the number of observations.

The dissociation was preserved in the same direction at every threshold, and no measure reversed sign across the three widths (Supplementary Fig. 7). The reward-learning differences indexed by the PALP were significant throughout: relative to ST, GT made more commission errors in the reward-reward (r=0.15, 0.18, 0.24 at the 10%, 20%, and 40% intermediate widths), reward-punishment (r=0.13, 0.15, 0.19), and punishment-punishment (r=0.09, 0.09, 0.14) conditions, and more omission errors in the reward-punishment (r=0.13, 0.14, 0.17) and reward-reward (r=0.14, 0.13, 0.20) conditions, and GT showed slower reaction times across all three conditions (reward-reward r=0.16, 0.18, 0.23; reward-punishment r=0.15, 0.17, 0.21; punishment-punishment r=0.10, 0.10, 0.15). The spatial working-memory advantage of ST (between-search errors r=0.15, 0.16, 0.21) and the higher CGT risk-adjustment in ST (r=0.11, 0.11, 0.18) were likewise significant at every threshold. A small number of additional measures (CGT deliberation time, punishment-punishment omission errors) reached significance only at the most restrictive 40% threshold, again in the direction observed in the primary analysis.

These results complement the k=2 analysis reported above (Supplementary Fig. 6). Under k=2, with the full intermediate band folded into the two groups, the reward-learning difference remains significant while the smaller working-memory and decision-making differences do not, whereas all of these differences are present when the intermediate band is set aside. This pattern is consistent with the intermediate participants being the less prototypical members of the two distributions.

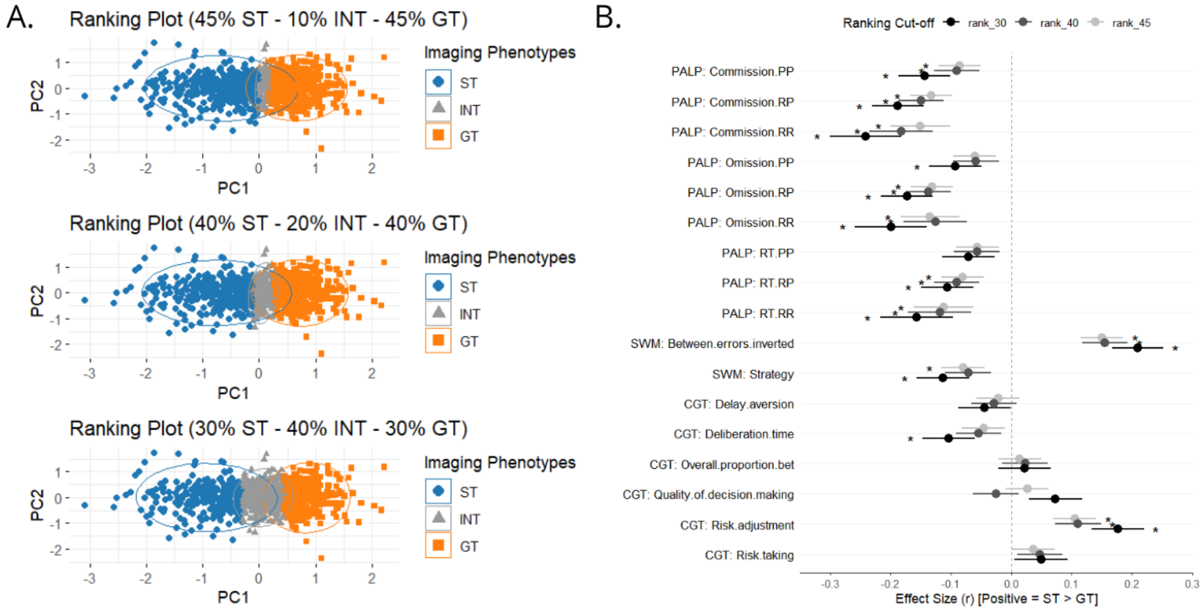


**Supplementary Fig. 7. Robustness of the ST/GT neuropsychological dissociation to the width of the intermediate group.** (A) Participants ranked on the first two principal components of the reward-related features, with the most ST-like and most GT-like 45%, 40%, and 30% classified as ST (blue) and GT (orange) and the remaining 10%, 20%, and 40% as intermediate (grey). (B) Effect size r of two-sided Wilcoxon rank-sum tests comparing ST and GT on each neuropsychological measure at each threshold (rank_45, rank_40, rank_30). Positive values indicate higher scores in ST. Stars denote $p < 0.05$.

#### Trial-by-trial analysis

The trial-by-trial analysis for the No-reward condition showed no differences in the ventral striatum activity between ST and GT (β=−0.0076, SE = 0.0276, t(245.3)=−0.276, p=0.783). The Trial effect in the ventral striatum responses indicates a slight decrease in BOLD activity over trials (β=−0.00181, SE = 0.00091, t(243.0)=−1.983, p=0.048), with no group differences in habituation rates (β=0.00160, SE = 0.00142, t(243.0)=1.126, p=0.261). In the Reward condition, ST showed significantly higher ventral striatum responses than GT (β=0.0860, SE = 0.0287, t(243.0)=2.990, p=0.003). There was no overall response decrease over trials ($\beta$=−0.00048, SE = 0.00091, t(241.9)=−0.526, p=0.599). However, a significant Trial × Group interaction ($\beta$=0.00275, SE = 0.00141, t(241.9)=2.540, p=0.008) indicated that ST maintained a stronger ventral striatum response over trials, compared to GT (Supplementary Fig. 8).

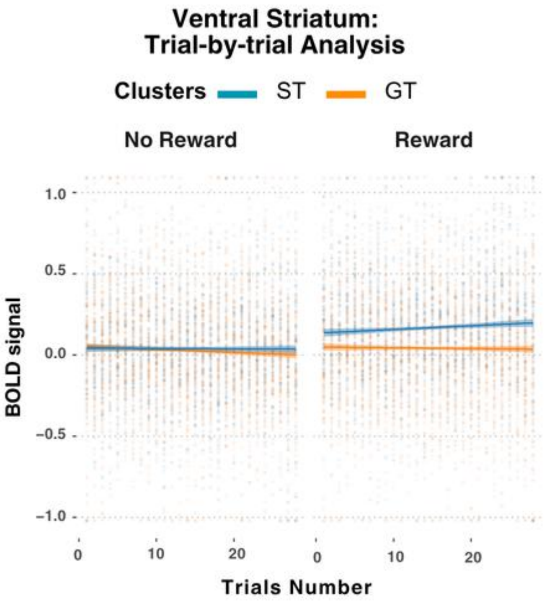


**Supplementary Fig. 8. Trial-by-trial analysis.** Trial-by-trial analysis showed ST had consistently higher ventral striatum activity than GT during reward trials, with no difference during no-reward trials.

#### Pharmacological Challenge cohorts

***Single-dose***

##### Dose-response effect of risperidone during the reward anticipation phase

The availability of risperidone conditions at increasing doses allowed us to investigate also dose–response effects, in this dataset (N=17), in particular in the anticipation phase, where a Cluster × Drug interaction had been reliably identified in the ventral striatum and the whole-network analysis, was further tested in the bilateral ventral striatum.

Results are shown in Supplementary Fig. 9, panel A. In ST, a significant overall Drug effect was observed ($\chi^2(2)$=10.829, p=0.005) with a robust linear trend ($\chi^2(1)$=10.800, p=0.001), indicating a dose-dependent decrease in BOLD response; the quadratic term was non-significant ($\chi^2(1)$=0.029, p=0.865). A stepwise comparison confirmed that high-dose risperidone significantly reduced ST responses relative to placebo and low-dose ($\chi^2(1)$=7.825, p=0.005). Post-hoc contrasts showed a significant difference between placebo and high-dose ($\beta$=–0.665, SE=0.181, t(14)=3.677, $p_{FDR}$=0.008), whereas the low-dose effect was not significant ($\beta$=–0.3075, SE=0.1808, t(14)=1.700, $p_{FDR}$=0.111).

For GT, no significant drug effects emerged ($\chi^2(2)$=1.978, p=0.372), with non-significant linear ($\chi^2(1)$=0.536, p=0.464) and quadratic ($\chi^2(1)$=1.442, p=0.230) trends. Post-hoc contrasts revealed no differences between placebo, low-dose, and high-dose (all $p_{FDR} \geq 0.532$). Bayesian analysis strongly supported the null hypothesis, showing no risperidone effect on anticipatory responses in GT (Bayes Factor, $BF_{01}$=5.47, 5.93, 3.95, respectively). The BF assesses the evidence that the data provide an effect of being present or absent. Specifically, "1" corresponds to H1 and "0" to H0. Generally, values of BF01 larger than 3 indicate evidence favoring H0 (i.e., no effect).

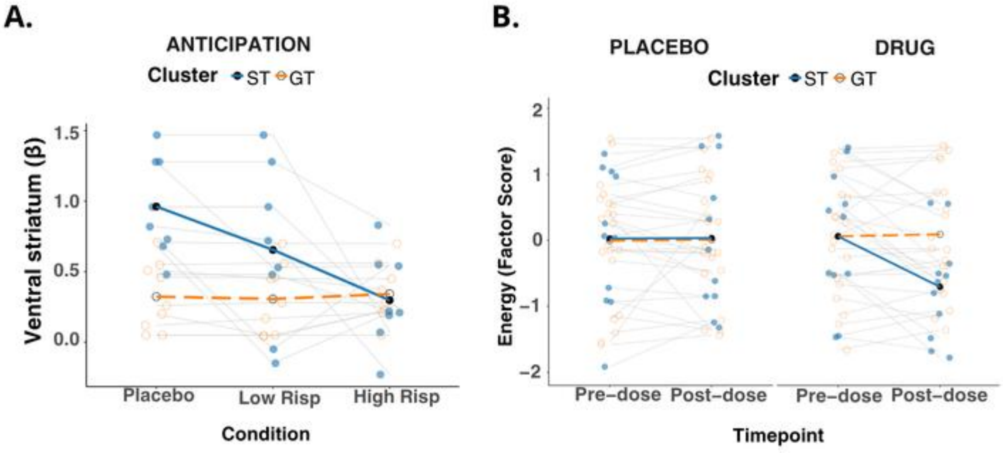


**Supplementary Fig. 9. Single-dose Pharmacological challenge cohort. (A)** Dose-response analyses indicated that ST anticipatory responses linearly decreased with increasing risperidone doses, while GT showed no change. **(B)** Behavioral analysis showed that ST had a lower Energy Factor Score after the drug dose. *Abbreviations*: ST=sign-trackers; GT=goal-trackers; BOLD=blood-oxygen-level-dependent.

##### Robustness of the pharmacological and clinical effects to classification and exclusion

Following the same logic applied to the Discovery and Replication cohorts, we tested whether the principal effects in the three smaller cohorts depended on the number of clusters used for projection or on the exclusion of non-concordant participants. In each cohort the principal effect was re-estimated under the labeling that span the available choices: the primary analysis reported in the manuscript, in which non-concordant participants are removed; projection of the Discovery centroids with all participants retained; and an independent within-sample $k = 2$ solution with all participants retained. For the single-dose cohort, projection was examined from both the three-group and the two-group Discovery solutions. Per-arm effects in the pharmacological cohorts were the simple main effects of the within-phase Drug × Cluster × Arm models, and clinical effects used the tests of the primary analysis. Full values are reported in Supplementary Table 4.

The principal effects were preserved in direction and significance under every labeling. In the single-dose cohort, the drug-related reduction of ST anticipatory ventral-striatal BOLD was significant whether non-concordant participants were removed or retained and whether labels were projected from the three-group or the two-group solution (drug − placebo from −0.41 to −0.51 SD, all $p \leq .0003$). In the repeated-dose cohort, amisulpride reduced ST anticipatory BOLD (−0.97 to −1.03 SD, all $p \leq .003$), and aripiprazole increased GT anticipatory BOLD (+0.86 to +0.90 SD, all $p \leq .023$) and reduced GT outcome BOLD (−0.93 to −1.08 SD, all $p \leq .021$), under every labeling. In the clinical cohort, the greater negative-symptom burden of GT relative to ST, and the correlations among D2 receptor affinity, anticipatory BOLD, and negative symptoms, remained significant whether non-concordant patients were removed or retained.

**Supplementary Tables**

**Supplementary Table 1. Demographic characteristics across the four study cohorts.** The table presents demographic data for all four cohorts, with the Discovery cohort further stratified by site. For each cohort and subgroup, the sample size, sex distribution (number of males and females), and mean age with standard deviation (SD) are reported. *Abbreviations:* M=males; F=females.

| Cohort | Sample Size (N) | M:F | Mean Age ± SD |
|---|---|---|---|
| ***Discovery*** | 890 | 436:454 | 22.07 ± 0.7 |
| **Sites** | | | |
| *Berlin* | 123 | 55:68 | 22.1 ± 0.6 |
| *Dresden* | 102 | 55:47 | 21.5 ± 0.7 |
| *Dublin* | 103 | 58:45 | 22.3 ± 0.7 |
| *Hamburg* | 122 | 62:60 | 22.1 ± 0.9 |
| *London* | 104 | 54:50 | 22.2 ± 0.6 |
| *Mannheim* | 115 | 55:60 | 22 ± 0.5 |
| *Nottingham* | 116 | 54:62 | 22.1 ± 0.6 |
| *Paris* | 105 | 43:62 | 22.2 ± 0.6 |
| ***Replication*** | 245 | 104:141 | 26 ± 6 |
| ***Pharmacological Challenge*** | | | |
| ***Single-dose*** | 34 | 34:0 | 26.9 ± 6.8 |
| ***Repeated-dose*** | 48 | 22:26 | 26.5 ± 8.15 |
| ***Clinical*** | 35 | 22:12 | 24.7 ± 5.6 |

**Supplementary Table 2. Whole-brain activation during the monetary incentive delay task in the Discovery, Replication, and Clinical cohorts.** Brain regions' activations were detected through one-sample t-tests on the Reward > No-Reward contrast during anticipation and outcome in the Discovery, Replication, and Clinical cohorts. All regions are shown at $p_{TFCE-FWE}<0.05$; cluster extent=20 voxels. *Abbreviations*: MNI=Montreal Neurological Institute; L=left; R= right; BA=Broadmann Area.

| Phase: Contrast | Group | MNI coordinates X Y Z | Regions |
|---|---|---|---|
| ***Discovery Cohort*** | | | |
| ***Anticipation: Reward > No Reward*** | *Sign-trackers* | -28 -78 -18 | Lingual Gyrus Fusiform L |
| | | -20 -78 -18 | BA18 |
| | | -36 -64 -20 | Cerebellum L |
| | | 10 -80 -14 | Lingual R BA18 |
| | | 12 -84 8 | Cuneus R |
| | | -30 -58 -22 | Cuneus R |
| | | 4 -86 6 | Calcarine R BA17 |
| | | 30 -76 -20 | BA19 Fusiform R |
| | | 12 -2 16 | Caudate Body R |
| | | -12 0 14 | Caudate L |
| ***Anticipation: Reward > No Reward*** | *Goal-trackers* | 10 -86 0 | Calcarine R |
| | | -16 -90 -12 | Lingual L |
| | | 38 -22 58 | BA4 Precentral R |
| | | 12 -16 8 | Thalamus R |
| | | -12 -18 8 | Thalamus L |
| | | -16 -24 12 | Pulvinar L |
| | | 14 16 -2 | Caudate R |
| | | -10 8 -2 | Pallidum L |
| | | 30 -54 -18 | Fusiform R |
| ***Outcome: Reward > No Reward*** | *Sign-trackers* | -6 2 24 | Corpus Callosum |
| | | 0 24 -6 | Anterior Cingulum L |
| | | -16 -26 -16 | ParaHippocampal L |
| | | -6 -32 -58 | Midbrain |
| | | 32 14 -18 | Insula R |
| ***Outcome: Reward > No Reward*** | *Goal-trackers* | 2 50 2 | Frontal Superior Medial R |
| | | 4 46 -6 | Frontal Medial Orbital R |
| | | 2 46 20 | Cingulum Anterior R |
| | | 0 60 4 | Frontal Superior Medial L |
| ***Replication Cohort*** | | | |
| ***Anticipation: Reward > No Reward*** | *Sign-trackers* | 4 0 58 | Supplementary Motor Area R |
| | | -4 6 56 | Supplementary Motor Area L |
| | | 28 16 -44 | Temporal Pole R |
| | | -42 14 54 | Primary Motor Cortex L |
| ***Anticipation:*** | *Goal-trackers* | | |

| | | | |
|---|---|---|---|
| ***Reward > No Reward*** | | | |
| ***Outcome: Reward > No Reward*** | *Sign-trackers* | -14 -88 -18 | Secondary Visual Cortex L |
| | | 2 -84 -10 | Secondary Visual Cortex R |
| | | -52 18 -10 | Superior Temporal Pole L |
| | | -40 18 -10 | Inferior Orbital Frontal Gyrus L |
| ***Outcome: Reward > No Reward*** | *Goal-trackers* | -16 -90 -6 | Middle Secondary Visual Cortex L |
| | | 14 -90 -2 | Primary Visual Cortex R |
| | | 14 -84 -12 | Secondary Visual Cortex R |
| ***Clinical cohort*** | | | |
| ***Anticipation: Reward > No Reward*** | *Sign-trackers* | -42 -24 58 | Supplementary Motor Area L |
| | | -38 -28 64 | Primary Motor Cortex L |
| | | -32 -40 58 | Primary Sensory Cortex L |
| | | 4 -14 44 | Ventral Anterior Cingulum R |
| | | -54 -22 34 | Supramarginal Gyrus L |
| | | -62 -18 18 | Primary Sensory L |
| | | 36 -16 18 | Insula R |
| | | -24 12 -8 | Putamen L |
| | | -14 -66 10 | Primary Visual Cortex R |
| | | 46 -12 54 | Primary Motor Cortex R |
| | | 58 -2 44 | Supplementary Motor Area R |
| ***Anticipation: Reward > No Reward*** | *Goal-trackers* | | |
| ***Outcome: Reward > No Reward*** | *Sign-trackers* | | |
| ***Outcome: Reward > No Reward*** | *Goal-trackers* | 54 26 34 | Dorsolateral Prefrontal Cortex R |
| | | 12 10 8 | Caudate R |
| | | -28 -96 12 | Secondary Visual Cortex L |
| | | 16 -62 8 | Primary Visual Cortex R |
| | | 50 -5 -18 | Superior Temporal Gyrus R |
| | | 24 -88 -8 | Secondary Visual Cortex R |
| | | -42 34 24 | Dorsolateral Prefrontal Cortex L |
| | | 22 -2 -20 | Amygdala R |
| | | -40 -12 12 | Insula L |
| | | -36 -30 16 | Primary Auditory L |
| | | 8 16 -12 | Subgenual R |
| | | -42 -12 18 | Primary Motor Cortex L |
| | | -50 -12 16 | Primary Sensory Cortex L |

**Supplementary Table 3. Descriptive statistics by imaging-derived clusters and Wilcoxon rank-sum test results in the Discovery and Replication cohorts.** This table reports descriptive and inferential statistics for each imaging-derived cluster across both the Discovery and Replication cohorts. Analyses were conducted on standardized data (scaled 0–1). Effect sizes (r equivalent) were computed using the *rstatix* R package [31], by dividing the Wilcoxon Z-transformed values by the square root of the number of observations [32]. For each variable, the table includes Wilcoxon Z-values, effect sizes, uncorrected p-values, and p-values corrected for multiple comparisons using the FDR correction ($p_{FDR} < 0.05$). Means and standard deviations (SD) are reported based on raw (non-standardized) data to facilitate interpretability. Cluster sample sizes (N) are also indicated. *Abbreviations:* FDR=False Discovery rate; ST=sign-trackers; GT=goal-trackers; SD=standard deviation; RT=reaction times, PP=punishment-punishment, RR=reward-reward, RP=reward-punishment.

| Sample | Questionnaire/Task | Score | Z | p(unc.) | $p_{FDR}$ | effect size r | Cluster ST Mean ± SD (N) | Cluster GT Mean ± SD (N) |
|---|---|---|---|---|---|---|---|---|
| **Discovery** | **Temperament and Character Inventory-Revised (TCI-R)** | *Disorderliness* | -1.902 | 0.057 | 0.135 | 0.087 | 21.65 ± 4.03 (160) | 21 ± 4.12 (317) |
| | | *Exploratory excitability* | -1.745 | 0.081 | 0.135 | 0.08 | 34.63 ± 4.24 (160) | 33.96 ± 4.24 (317) |
| | | *Extravagance* | -1.777 | 0.076 | 0.135 | 0.081 | 26.21 ± 3.69 (160) | 26.78 ± 4.03 (317) |
| | | *Impulsiveness* | -0.71 | 0.478 | 0.478 | 0.033 | 25.27 ± 3.96 (160) | 24.96 ± 4.15 (317) |
| | | *Novelty Seeking* | -1.411 | 0.158 | 0.198 | 0.065 | 107.78 ± 9.18 (160) | 106.70 ± 10.46 (317) |
| | **Substance Use Risk Profile Scale (SURPS)** | *Anxiety sensitivity* | -1.635 | 0.102 | 0.379 | 0.075 | 11.31 ± 2.51 (160) | 11.75 ± 2.51(317) |
| | | *Hopelessness* | -0.624 | 0.533 | 0.710 | 0.029 | 12.63 ± 3.5 (160) | 12.89 ± 3.77 (317) |
| | | *Impulsivity* | -0.289 | 0.773 | 0.773 | 0.013 | 10.67 ± 2.11 (160) | 10.7 ± 2.27 (317) |
| | | *Sensation seeking* | -1.313 | 0.189 | 0.379 | 0.06 | 16.84 ± 3.37 (160) | 16.47 ± 3.38 (317) |
| | **Cambridge guessing task (CGT)** | *Delay aversion* | -0.173 | 0.863 | 0.863 | 0.01 | 0.165 ± 0.13 (156) | 0.17 ± 0.15 (314) |
| | | *Deliberation time* | -1.502 | 0.133 | 0.399 | 0.07 | 1510.02 ± 443.43 (156) | 1569.68± 478.33 (314) |
| | | *Overall proportion bet* | -0.322 | 0.748 | 0.863 | 0.02 | 0.51 ± 0.13 (156) | 0.51± 0.13 (314) |

| | | | | | | | | |
|---|---|---|---|---|---|---|---|---|
| | | *Quality of decision making* | -0.662 | 0.508 | 0.863 | 0.03 | 0.97 ± 0.05 (156) | 0.97 ± 0.05 (314) |
| | | *Risk adjustment* | -3.588 | 0.000 | 0.002 | 0.17 | 2.25 ± 0.93 (156) | 1.93 ± 1.03 (314) |
| | | *Risk taking* | -0.299 | 0.765 | 0.863 | 0.01 | 0.57 ± 0.12 (156) | 0.57 ± 0.13 (314) |
| | **Passive Avoidance Learning Paradigm (PALP)** | *Commission RP* | -3.040 | 0.000 | 0.001 | 0.14 | 0.078 ± 0.07 (158) | 0.11 ± 0.09 (315) |
| | | *Commission PP* | -3.945 | 0.002 | 0.005 | 0.18 | 0.08 ± 0.08 (160) | 0.11 ± 0.1 (315) |
| | | *Commission RR* | -3.713 | 0.000 | 0.001 | 0.24 | 0.06 ± 0.08 (81) | 0.1 ± 0.09 (162) |
| | | *Omission PP* | -2.449 | 0.014 | 0.018 | 0.11 | 0.03 ± 0.04 (160) | 0.03 ± 0.05 (315) |
| | | *Omission RP* | -3.394 | 0.001 | 0.002 | 0.16 | 0.02 ± 0.05 (158) | 0.05 ± 0.05 (315) |
| | | *Omission RR* | -2.530 | 0.011 | 0.017 | 0.16 | 0.02 ± 0.04 (81) | 0.03 ± 0.04 (162) |
| | | *RT PP (ms)* | -1.277 | 0.202 | 0.202 | 0.06 | 876.58 ± 206.03 (160) | 905.66 ± 221.4 (315) |
| | | *RT RP (ms)* | -2.743 | 0.006 | 0.011 | 0.13 | 820.57 ± 173.99 (158) | 876.11 ± 192.67 (315) |
| | | *RT RR (ms)* | -1.825 | 0.068 | 0.077 | 0.12 | 857.91 ± 213.33 (81) | 918.65 ± 255.19 (162) |
| **Replication** | **Temperament and Character Inventory (TCI)** | *Cooperativeness* | -0.962 | 0.336 | 0.456 | 0.08 | 19.73 ± 3.6 (42) | 19.08 ± 4.01 (110) |
| | | *Harm Avoidance* | -1.05 | 0.294 | 0.456 | 0.09 | 8.96 ± 4.11 (42) | 8.4 ± 4.06 (110) |
| | | *Novelty Seeking* | -0.93 | 0.352 | 0.456 | 0.08 | 7.89 ± 4.52 (42) | 8.5 ± 4.43 (110) |
| | | *Persistance* | -0.336 | 0.737 | 0.737 | 0.03 | 3.05 ± 1.56 (42) | 2.97 ± 1.91 (110) |
| | | *Reward Dependence* | -0.859 | 0.390 | 0.456 | 0.07 | 9.7 ± 2.43 (42) | 9.4 ± 2.47 (110) |
| | | *Self Directedness* | -1.03 | 0.303 | 0.456 | 0.08 | 19.626 ± 6.01 (42) | 18.62 ± 6.14 (110) |
| | | *Self-Trascendeces* | -0.903 | 0.366 | 0.456 | 0.07 | 5.3 ± 6.25 (42) | 6.01 ± 6.36 (110) |

| | | | | | | | | |
|---|---|---|---|---|---|---|---|---|
| | **Wisconsin Card Sorting Test (WCST)** | *Number of Completed Categories* | -1.144 | 0.126 | 0.126 | 0.09 | 5.86 ± 0.53 (51) | 5.67 ± 1.06 (129) |
| | | *Number of Non Perseverative Errors* | -2.019 | 0.022 | 0.042 | 0.15 | 7.06 ± 7.03 (51) | 9.58 ± 9.35 (127) |
| | | *Number of Perseverative Errors* | -1.861 | 0.031 | 0.042 | 0.14 | 7.06 ± 3.75 (51) | 9.95 ± 8.46 (128) |
| | | *Number of trials* | -2.484 | 0.007 | 0.026 | 0.19 | 80.54 ± 15.84 (51) | 88.33 ± 20.42 (128) |
| | **Continuous Performance Test (CPT-AX)** | *HIT* | -0.143 | 0.886 | 0.430 | 0.01 | 0.97 ± 0.04 (50) | 0.98 ± 0.09 (128) |
| | | *FA* | -0.801 | 0.423 | 0.430 | 0.06 | 0.06 ± 0.06 (50) | 0.06 ± 0.07 (128) |
| | | *FAC* | -1.072 | 0.284 | 0.662 | 0.08 | 0.02± 0.03 (50) | 0.04 ± 0.06 (128) |
| | | *AP S* | -0.115 | 0.909 | 0.740 | 0.01 | 0.97 ± 0.03 (50) | 0.98 ± 0.03 (128) |
| | | *AY* | -1.826 | 0.068 | 0.909 | 0.14 | 0.09 ± 0.11 (50) | 0.08 ± 0.1 (128) |
| | | *DP S* | -0.199 | 0.842 | 0.909 | 0.02 | 4.19 ± 0.99 (50) | 4.18 ± 1.04 (128) |
| | | *DPC S* | -1.543 | 0.123 | 0.909 | 0.12 | 4.91 ± 0.97 (50) | 4.62 ± 1.11 (128) |

**Supplementary Table 4. Robustness of the pharmacological and clinical effects to clustering and exclusion.** For each cohort (single-dose risperidone/haloperidol, repeated-dose amisulpride and aripiprazole, and clinical) every effect is reported under three classifications. The first is the primary analysis used in the manuscript, in which the three-group Discovery solution is projected onto the cohort and non-concordant individuals are removed. The second projects the two-group Discovery solution onto the cohort and retains every individual. The third derives an independent within-cohort k = 2 solution and likewise retains every individual. Columns give the effect or analysis, the classification or sample, the sample size (with ST/GT counts shown where applicable), the effect size or test statistic, and the p-value. Effect size is the drug minus placebo contrast in SD units for the pharmacological rows, the partial correlation controlling age and sex for the clinical brain-behavior relationships, and the Mann-Whitney statistic for the clinical group contrast. Bold p-values indicate $p < .05$.

| Effect / analysis | Classification / sample | N (ST/GT) | Effect size / statistic | p |
|---|---|---|---|---|
| **Single-dose — risperidone/haloperidol (D2/D3 antagonist): ventral striatum, drug − placebo** | | | | |
| ST anticipation (reduced) | Primary (manuscript; non-concordant removed) | 32 (11/21) | −0.41 | **<.001** |
| | k=2 projection (all retained) | 34 (12/22) | −0.51 | **.0002** |
| | within-sample k=2 (all retained) | 34 (11/23) | −0.51 | **.0003** |
| **Repeated-dose — amisulpride (D2/D3 antagonist): ventral striatum, drug − placebo** | | | | |
| ST anticipation (reduced) | Primary (concordant; non-concordant removed) | 18 (13/5) | −0.98 | **.003** |
| | k=2 projection (all retained) | 23 (13/10) | −1.03 | **.002** |
| | within-sample k=2 (all retained) | 23 (18/5) | −0.97 | **<.001** |
| **Repeated-dose — aripiprazole (D2/D3 partial agonist): ventral striatum, drug − placebo** | | | | |
| GT anticipation (increased) | Primary (concordant; non-concordant removed) | 22 (13/9) | +0.86 | **.023** |
| | k=2 projection (all retained) | 25 (13/12) | +0.88 | **.008** |
| | within-sample k=2 (all retained) | 25 (16/9) | +0.90 | **.020** |
| GT outcome (reduced) | Primary (concordant; non-concordant removed) | 22 (13/9) | −1.07 | **.021** |
| | k=2 projection (all retained) | 25 (13/12) | −0.93 | **.020** |
| | within-sample k=2 (all retained) | 25 (16/9) | −1.08 | **.018** |
| **Clinical cohort** | | | | |
| GT > ST negative symptoms | Primary analysis (manuscript) | 25 (14/11) | W = 144 | **.006** |
| | Projection, all retained | 33 (14/19) | U = 210 | **.005** |
| | Within-cohort k = 2, all retained | 33 (22/11) | U = 212 | **<.001** |
| D2 affinity ~ anticipatory BOLD | Primary analysis (manuscript) | 24 | r = −0.46 | **.023** |
| | All patients retained | 30 | r = −0.47 | **.011** |
| Anticipatory BOLD ~ negative symptoms | Primary analysis (manuscript) | 25 | r = −0.58 | **.002** |
| | All patients retained | 33 | r = −0.49 | **.005** |
| D2 affinity ~ negative symptoms | Primary analysis (manuscript) | 24 | r = +0.47 | **.020** |
| | All patients retained | 30 | r = +0.46 | **.014** |
